# Spiking burstiness and working memory in the human medial temporal lobe

**DOI:** 10.1101/2022.08.08.503153

**Authors:** Francesco Cocina, Andreas Vitalis, Amedeo Caflisch

## Abstract

Persistent activity has commonly been considered to be a hallmark of working memory (WM). Recent evidence indicates that neuronal discharges in the medial temporal lobe (MTL) are compatible with WM neural patterns observed in cortical areas. However, the characterization of this activity rarely consists of measurements other than firing rates of single neurons. Moreover, a varied repertoire of firing dynamics has been reported in the MTL regions, which motivate the more detailed examination of the relationships between WM processes and discharge patterns undertaken here. Specifically, we investigate at different resolution levels firing irregularities in electrode recordings from the hippocampus, amygdala, and the entorhinal cortex of epileptic patients during a WM task. We show that some types of (ir)regularities predict response times of the patients depending on the trial periods under consideration. Prominent burst activity at the population level is observed in the amygdala and entorhinal cortex during memory retrieval. In general, regular and bursty neurons contribute to the decoding of the memory load, yet they display important differences across the three anatomical areas. Our results suggest that non-random (non-Poisson) patterns are relevant for WM, which calls for the development and use of statistics complementary to mere spike counts.

## 1 Introduction

A great deal of work has been dedicated over the last 50 years to the identification and characterization of neural activity observed during working memory (WM) tasks (Fuster and Alexander, 1971; Wang, 2021). The ability to maintain information in memory for a limited period of time has been traditionally explained in terms of biological mechanism by the presence of persistent spiking activity during the maintenance, or delay, period of the task. The term ‘persistent’ has been used in association with the elevated spike count of a restricted subset of neurons (Wang, 2021; Leavitt et al., 2017; Constantinidis et al., 2018) as well as to describe low-dimensional neural trajectories residing in an attractor state (Masse et al., 2020; Kamiński and Rutishauser, 2020). The brain areas typically associated to WM are the prefrontal, parietal, and sensory cortex (Christophel et al., 2017). However, recently also the medial temporal lobe (MTL) has attracted attention following the identification of signatures of persistent activity (Kamiński et al., 2017; Kornblith et al., 2017; Bausch et al., 2021) (see also earlier work Axmacher et al. (2007)).

Despite the multiple reports, a precise characterization of the persistent activity in terms of firing dynamic is missing. In addition, the properties of this activity that allow for an efficient coding of the memory variables are not fully understood. Here, we ask the question if bursty neurons and/or population bursts contribute to the decoding of WM signals. To answer this question, we investigate firing irregularities in the MTL during a modified Sternberg WM task. We utilize a data set of recordings from a region comprising the hippocampus, amygdala, and the entorhinal cortex of epileptic patients (available online (Boran et al., 2019, 2020)). In the original work (Boran et al., 2019), the authors hypothesized the involvement of the hippocampus in the WM process, related in particular to the memory load, and they used sets of 4, 6, or 8 letters written on a screen to be memorized. These authors found two sets of neurons with significantly enhanced activity during the delay phase and the memory retrieval phase, which were termed maintenance and probe neurons, respectively. An attractor-driven dynamics was postulated for the maintenance of WM information at the population level.

Higher irregularities were previously observed in the prefrontal cortex during the delay period of a delayed response task (Compte et al., 2003). Furthermore, large deviations in inter-spike interval (ISI) statistics from a Poisson process were unveiled across the cortex with differential patterns from sensory-motor to higher cortical regions (Maimon and Assad, 2009). In the human MTL, long-range temporal correlations among the ISIs were observed in both amygdala and hippocampus in spontaneous activity (Bhattacharya et al., 2005). More recently, a wealth of distinct irregular patterns associated with temporal coding during different memory phases was identified in the hippocampus and entorhinal cortex (Umbach et al., 2020).

In contrast to the original analysis, we here do not focus exclusively on the firing rates of single neurons but rather investigate the discharge patterns in terms of burstiness, i.e., irregularity of the spike sequences. In particular, we analyse the spike trains by adopting, predominantly, a local variation metric (Shinomoto et al., 2003, 2009) for the quantification of irregularities, which is independent from raw firing rates as we demonstrate below. Our analysis is structured as follows: first, we validate the local variation metric and investigate if it can be reliably related to trial and behavioural variables; second, we examine the burst activity at the population level and assess how it reports on memory processes and on the coordination of single units; a Fano factor analysis then helps clarify some of the previous results. Eventually, we study how the population decoding of trial variables depends on the firing behaviour of the underlying units. Our analysis sheds light on intrinsic differences between the dynamics of the three anatomical areas. It also reveals the relevance of non-random (non-Poisson) spike trains for memory maintenance and behavioural performance, i.e., response times.

## 2 Materials and Methods

### 2.1 Experimental design and recordings

Detailed information about subjects, task, recording setup, and spike sorting procedure can be found in Boran et al. (2020). All subjects provided written informed consent for the study. In this section, we recapitulate the aspects that strictly concern our analysis and describe the data filtering procedure. Subjects are asked to perform a modified Sternberg task, which comprises epochs of encoding, maintenance, and recall of memory items. In detail, after an initial period of fixating a blank screen (1 s), a stimulus is presented. This consists of a set of central four, six, or eight letters, possibly surrounded by ‘X’s on both sides such that the number of characters is always equal to eight. The letter ‘X’ is never part of the set to be memorized. The encoding period lasts for two seconds and is followed by a maintenance, or delay, period of three seconds where a blank screen is shown again. A probe letter is eventually presented and the subjects are instructed to answer rapidly whether the letter does or does not belong to the stimulus set (IN or OUT button press). The set sizes are randomly selected, except when the subject response is wrong. In that case, the set size of the subsequent trial is always chosen as four. A session is composed of 50 trials and lasts approximately 10 minutes.

Depth electrodes were implanted in the MTL of epileptic patients for potential surgical resection of their seizure foci. Intracranial electroencephalography (iEEG) recordings were performed into hippocampus, amygdala and the entorhinal cortex with depth electrodes combining macro- and micro-contacts (1.3 mm diameter, 8 macro-contacts of 1.6 mm length, spacing between contact centers 5mm, nine micro-contacts protruding radially 4mm from its tip, ADTech^®^). All of the trial recordings have a fixed length of 8 seconds and they all start with the fixation period (1 s). This implies that when response times are longer than 2 secs a part of the neural recordings preceding the response is missing. In many cases the trials affected are discarded; this is properly noted where required. Trials containing artifacts were also not considered throughout the analysis (Boran et al., 2019).

Spike sorting has been performed through the Combinato package (Niediek et al., 2016), and its results are provided with the data set. In our analysis, only neurons with an average firing rate > 1 Hz across trials were kept. Also, to control for potential cross-talk between electrodes, Jaccard similarity was computed between binarized spike trains (1 ms bin) of simultaneously recorded units. All of the session trials were concatenated for the calculation. Values of Jaccard similarity higher than 0.3 were considered suspicious, and sequentially selected units were discarded until all values fell below this threshold. In detail, we identified the unit that was contributing to the highest number of Jaccard values > 0.3 and removed it. If two units were equally contributing, we discarded the one with lower firing rate. The procedure was repeated until the threshold criterion was fulfilled globally. In the end, 7 out of 26 sessions were affected with a total number of discarded units equal to 35 (out of 992 previously selected).

### 2.2 Firing irregularities

#### *LvR* metric

Spiking irregularities were investigated by examining the sequence of the ISIs. We adopted an enhanced local variation measure (*LvR*) to quantify the firing (ir)regularities of the units during single trials (Shinomoto et al., 2003, 2009). Unlike other common metrics, such as the coefficient of variation (CV), *LvR* accounts for fluctuations in firing rates along the time series and, also, corrects for the refractory period following a spike. It is computed as

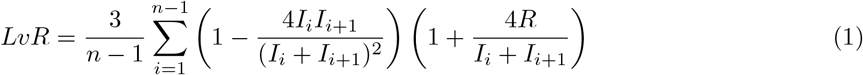

where *I* indicates the ISI, *n* the total number of ISIs, and *R* the refractoriness constant. This latter parameter is set to 5 ms (as in the original work of Shinomoto et al. (2009)) for the single-unit calculations. For the combination of two or more units, the refractoriness correction was not considered (*R* = 0). *LvR* values were computed for (windows of) spike sequences containing at least five spikes. When this condition was not met, the corresponding data points were simply discarded unless noted otherwise.

#### Change points (CPs)

We identified sharp variations in the single-unit firing rates during each trial through an adaptive CP procedure (Gallistel et al., 2004; Jezzini et al., 2013). The method uses the empirical cumulative count of spikes and compares it with the expected one, which, in our case, is the one deriving from a perfectly regular firing with a matched number of spikes (*i*.*e*., a uniform distribution).The earliest time point where the two distributions differ maximally is considered and identified as a CP contingent upon the result of a binomial test between the spike counts before and after that point (see Gallistel et al. (2004) for more details). After a CP is evaluated to be significant, the algorithm is applied again to the remaining data following the CP. The adaptive element of the procedure rests in reducing progressively the *p* value confidence threshold until no change points are identified during the fixation period (Jezzini et al., 2013). This is done in practice by increasing the 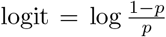 in steps of 0.2 from 1.3 to 5.9 (corresponding to a range *p* ≃ 0.05 − 10^−6^).

### 2.3 Time-resolved analysis of irregularities and trial variables

To investigate the relation between *LvR* and response times, we calculated the Pearson correlations between these two quantities using data from the whole pool of sessions and trials (Fig. 4). Differently, in order to assess whether there are significant regions in the *LvR* spectrum tuned to a particular trial class, *i*.*e*., set size or response correctness (Fig. S3), we devised the following test. The *LvR* values are first collected across all of the sessions and trials as above and then rank-transformed in ascending order. This was done separately for single as well as combined (2 and 3 units, see Fig. 3a) *LvR* values. For the combined case, we subsampled only a fraction of points (see below). The cumulative distributions of the rank-transformed *LvR* values are then computed for the single classes, *e*.*g*., correct and wrong responses. The rank transformation makes the (following) statistic invariant to the underlying distributions (see Fig. S2a) and guarantees a fair comparison between the different combinations. We took the sum of the point-wise differences between the two curves as the test statistic for quantifying the dissimilarity between the two distributions. For the set size grouping variable, the difference was between high (6 and 8) and low workloads (4); for the response correctness, we considered the difference between the wrong and correct subsets. For this latter grouping variable, we considered only sessions with at least five wrong trials. The statistic obtained is equal to the total area between the two curves but respecting the sign of the difference. It differs from the more known Wasserstein, or earth-mover’s, distance as there the absolute difference between the curves is considered.

Next, we describe the procedures for subsampling the data set and calculating the chance level distribution which are conceptually the same also for both of the analyses introduced before. In order to maintain across the different unit combinations (single, 2, or 3) the same number of points used for calculating the test statistic (the Pearson correlation or the dissimilarity between the class distributions), we subsampled, within each session, from the combined *LvR* values (2 and 3) a number of points equal to the ones in the single-unit calculation. From this, the test statistic is computed, and the whole process is repeated 100 times. Within each repetition (and also for the single-unit *LvR*), we generated a null distribution by permuting 1000 times the trial variables, *i*.*e*., response times, set sizes, or correct/wrong responses. This permutation was performed preserving both the trial structure and the subject identity, that is, units that were simultaneously recorded were assigned the same trial variable, drawn from the ones available for that specific subject. By comparing the true statistic value to the null distribution, a *p* value could be extracted. The summary *p* value for the combined *LvR* is given by the median of the 100 extracted ones.

### 2.4 Fano factor

Mean-matched Fano factors (FF) were calculated to investigate the trial variability of the spiking responses (Churchland et al., 2010). In contrast to the raw FF calculation, that is, the simple ratio between variance and mean of the spike counts, the mean-matched procedure controls for local variations in firing rates that can trivially affect the FF calculation. For example, similar to *LvR*, the statistic should account for the fact that refractoriness periods might reduce the spiking variability in periods of high firing rates, and as a consequence, also across trials, leading to an artificial decrease. The calculation starts by extracting mean and variance of the spike counts for each combination of units and conditions (here, the three set sizes). We adopted sliding windows of 500 ms with a time step of 50 ms. The total numbers of points/combinations were 1254, 699, 919 for hippocampus, amygdala, and entorhinal cortex, respectively. The greatest common distribution of mean spike counts across all the time windows was extracted (in practice, a histogram with bin size of 0.5 was employed). For each window, we discarded points randomly such that the common distribution was matched and, eventually, FF was determined as the slope of the line regressing variance over the mean of the remaining points. For the linear regression the intercept was constrained to zero, and each point was weighted by the (inverse of 0.01 plus) standard error of the variance. Due to the multiple mean-matching possibilities, the procedure was repeated 50 times and the average FF with 95% confidence intervals (CI) was reported.

### 2.5 Population bursts

A population burst is defined as a period of sustained collective activity where the population firing rate exceeds a certain threshold for at least 100 ms. Our procedure closely resembles the one in Vaz et al. (2020). Instantaneous firing rates were extracted by convolving the spike rasters of each unit with a Gaussian kernel of bandwidth equal to 25 ms and subsampling with a 10-ms step. These time series were used for calculating both the population firing rate, as the average value across the units, and the threshold, which is described in the following. In turn, (a) single-unit firing rates were averaged over the whole trial window; (b) these resulting values were then averaged across units within each trial; (c) the mean and standard deviation across the trials were extracted. These last two values eventually served to define the threshold as mean + 3 · s.d. If two consecutive bursts were closer than 150 ms (tail to head), the one with lower firing rate was discarded. We assigned the single burst events to a specific trial period if at least 80% of the burst window was located within its boundaries. When dealing with the probe period, the right boundary was always defined by the response time.

The unit composition of a single burst, ***w***, was defined as a vector composed by the averages of the single-unit firing rates within the burst window. The smoothed firing rates (25-ms Gaussian kernel) were again employed.

The sparsity measure quantifies the prevalence of a group of units in the population burst activity. It is computed as follows:

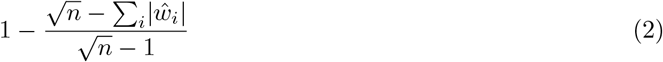

where *n* is equal to the number of units (the length of ***w***), and 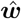 is the unit composition scaled to unit length. Sparsity values close to zero indicate the net prevalence of few units to the burst activity whereas higher values suggest a more balanced contribution of all of the units. The *LvR* of the burst events was calculated as the weighted mean of the single-unit *LvR* using the unit composition elements ***w*** as weights. To quantify the variability of the composition of bursts across the different trials, we adopted the following procedure. Within each trial, we computed the average composition ***w*** of the bursts present therein and normalized it to unit length 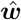. Subsequently, we computed the Euclidean distances of these vectors between all of the different trials where bursts were present. The mean (*µ*_*B*_) and variance 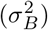 of this distance matrix (only the upper triangle) were calculated. While *µ*_*B*_ and 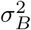 describe the variability between burst compositions, we chose to derive a normalized measure from them, introducing a reference value. Specifically, we applied the same procedure described above to the firing rates computed within the entire trial (*i*.*e*., not only within windows corresponding to bursts), thus obtaining mean *µ*_*T*_ and variance 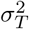. The final variability measure for burst composition was then chosen as a d-prime, or sensibility index, as follows:

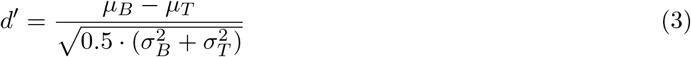

To test differences between set sizes, the Mahalanobis distance was calculated between single bursts and those of a reference group - in this case those in the fixation period - by using the unit composition of the bursts as the metric space. In order to account for the intrinsic (*i*.*e*., workload-blind and trial-wise) variability of the burst composition, we properly rescaled each unit dimension such that high firing rates, with higher deviations, do not dominate the calculation. The distance was computed as follows:

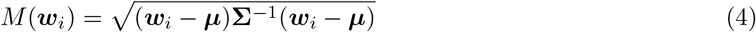

where ***µ*** and **Σ** represent the mean vector and the covariance matrix, respectively, extracted from the collection of all of the fixation bursts. For each anatomical region we selected those sessions with at least five recorded units and ten bursts overall during the fixation periods. Eventually, we averaged across the single *M* (***w***_*i*_) within each combination of trial period and set size.

### 2.6 Decoding analysis

Decoding analysis was performed on the neuronal pseudo-population obtained by collecting units across all of the sessions and patients. Spike counts were computed per unit in non-overlapping windows of 250 ms within the 3 s maintenance period, thus yielding 12 points per trial. A *z* score normalization was then applied to each of the neuronal series. No smoothing was applied. The order and number of each grouping variable, *e*.*g*., set size, differ between each session, thus preventing an alignment of the trials of different sessions based solely on temporal succession. To this end, we describe in the following a bootstrap procedure adopted to calculate the decoding accuracies (Meyers et al., 2008). First, in each session we identified the minimum between the number of trials belonging to each of the classes. If that number was lower than five, that session was discarded. The minimum of these values across all of the session, *n*_*T*_, was taken as the number of trials to be sampled per class within each session. Following this criterion, the classes’ instances were thus forcefully balanced. Second, after sampling the trials, the resulting data sets from each session were concatenated together creating the final pseudo-population activity matrix. The number of data points is equal to 12 × 2 × *n*_*T*_ given that we perform only binary classifications (*n*_*T*_ = 9, set sizes 4 and 6-8; *n*_*T*_ = 6, correct and wrong responses). At last, we train and test a linear support vector machine (SVM) following a ten-fold cross-validation scheme (SVC in Python package *scikit-learn*). The mean accuracy score across the ten test sets was computed. The entire procedure was repeated 50 times, sampling thus different alignment configurations between trials of different sessions. To generate a null distribution of accuracy scores, we shuffled the class labels 500 times within each bootstrap cycle and repeated the procedure described above. One-tailed *p* values were extracted and the median of these 50 quantities was taken as the summary value.

The total numbers of units for hippocampus, amygdala, and entorhinal cortex utilized when discriminating the set size were 418, 233, and 308, respectively; when classifying the response correctness, the numbers of units were correspondingly 103, 66, and 58. A large part of the decoding analysis is performed on subsets of these units defined by the *LvR* values. The neuronal pseudo-population is split into n-tiles according to the mean *LvR* value computed over the trials composing each particular bootstrap cycle. This implies that the same unit in distinct cycles can be assigned to different n-tiles.

To quantify the variation in accuracy when removing a n-tile with respect to the full population, we computed within each bootstrap cycle a d-prime index as follows:

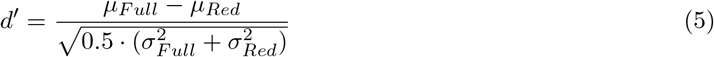

where the subscripts indicate the full or the reduced neuronal population, and *µ* and *σ*^2^ are, respectively, the mean and variance of the accuracy scores across the ten test sets. We ensured that the classification procedures of the two populations were performed on the same sample of trials. Eventually, we computed mean and SEM of the *d*′ indexes and performed a *t* test with zero as null hypothesis for the sample mean.

### 2.7 Statistical analysis

All statistical tests were two-tailed unless stated otherwise. The utilization of non-parametric tests (Kruskal-Wallis, Wilcoxon rank sum, Wilcoxon signed rank tests) over parametric ones (ANOVA, *t* tests) in one-way comparisons was decided upon significance of at least one Shapiro-Wilk test of normality on the samples involved (*p* < 0.05). Post-hoc, pairwise tests were adjusted for multiple comparison with the Benjamini-Hochberg (BH) procedure. This multiple comparison adjustment is also the one applied in general where noted.

### 2.8 Code and software accessibility

The analyses were carried out with R and Python packages available online. Customized code and scripts supporting the current study will be uploaded to https://gitlab.com/CaflischLab.

## 3 Results

The experiment is presented in Sec. 2.1 (see also Fig. 1a and the description in the published data set (Boran et al., 2020)). In the following we will use also the term ‘neuron’ to indicate a unit.

**Figure 1:**
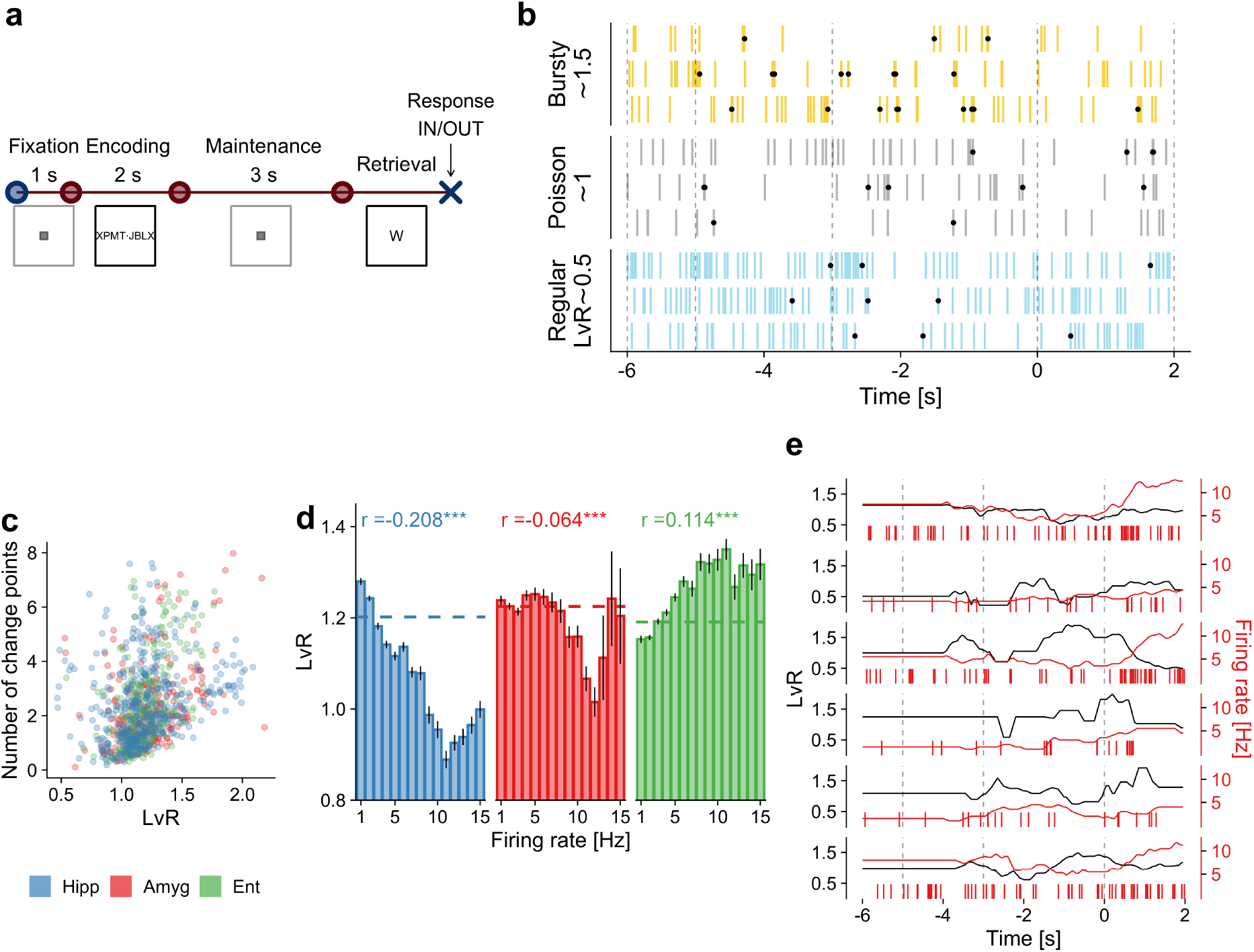
Memory task and firing irregularity of single neurons. (a) A modified Sternberg WM task was performed by epileptic patients. In each trial a set of for, six, or eight letters was displayed on a screen during the encoding period (2 s). A delay period followed (3 s; maintenance) and, after that, a probe letter appeared on the screen (retrieval). The subject then had to indicate whether the letter belonged to the set. (b) Examples of different firing patterns. The three types of firing characterizations shown derive from *LvR* values distributed around 0.5, 1, and 1.5 for, respectively, regular, Poisson, and bursty dynamics. Dots indicate change points (CPs). (c) Scatter plot between *LvR* and number of CPs. The two quantities are computed for each unit by averaging over the single trials. (d) Relationship between burstiness and firing rates. Firing rates computed for each neuron (418, 233, 306 units for hippocampus, amygdala, and entorhinal cortex, respectively) were grouped into 1-Hz bins. Mean *LvR* values with SEM are shown for each bin. Dashed lines indicate the average *LvR* values. Pearson correlation value between *LvR* and firing rates computed over the whole neuronal population are shown on top. (e) Examples of time series of *LvR* and firing rates values. Both of the quantities are calculated on a sliding window of 2-s with 50-ms steps. Windows are right-aligned to the time index (hence, flat values in the first two seconds). Burstiness levels for the population of maintenance and probe neurons (Boran et al., 2019) and trial-to-trial variability of the *LvR* values are examined in Fig. S1.

We start by investigating the firing irregularities, or burstiness, of the single units by using a metric that relies on the interspike intervals (ISIs). We adopt a coefficient of local variation while correcting for refractoriness defined as *LvR* (Shinomoto et al., 2003, 2009) (see 2.2 for details). The *LvR* metric takes into account the local variations of consecutive ISIs, and its value characterizes the spike train dynamic as regular (LvR < 1), random Poisson-distributed (∼1), or bursty (> 1), as depicted in Fig. 1b.

In order to assess the validity of this metric, we used another proxy measure for quantifying irregularities in the spike trains, namely, the number of change points (CP). Change points are designed to locate and indicate sudden variations in firing rates. We hypothesized that a higher count of CPs will be observed for more irregular, burstier dynamics (see Fig. 1b as an example, see Sec. 2.2 for details). The number of CPs consistently exhibits significant correlations with *LvR* in all of the anatomical areas (*r* = 0.20, *t*(17626) = 26.5; *r* = 0.24, *t*(9747) = 24.8; *r* = 0.26, *t*(13102) = 30.5 for hippocampus, amygdala, and entorhinal cortex, respectively; all *p* values < 10^−10^, Student’s *t* test) (Fig. 1c). Notably, the CP count does not fulfill our requirement of being independent of firing rates (correlations of *r* = 0.38, 0.46, 0.36, respectively, as before; all *p* values < 0.001), thus highlighting that the *LvR* metric is a suitable approach to the nontrivial task of unveiling results that depend specifically on actual firing (ir)regularities rather than just rates.

### Irregularities show a non-trivial relationship with firing rates

By construction, *LvR* is invariant to gradual firing rate fluctuations along time series, and, importantly, it does not depend on differences in spike counts between units (Shinomoto et al., 2009). In the following, we test the hypothesis that *LvR* and firing rates have no trivial interdependence explicitly and in more depth. This is crucial for the remainder of the analysis as we desire to obtain information that is complementary to that provided by the firing rates for describing the spike sequences. We start in Fig. 1d by showing the average *LvR* values for single bins of firing rate (1 Hz). In all of the three anatomical areas, the *LvR* trend for increasing firing rates is neither linear nor strictly monotonic. Moreover, although some strong correlations appear in specific intervals of firing rates, *e*.*g*., in the hippocampus from 1 to 10 Hz, these dependencies are not generally replicated in the other anatomical areas. The diverse relationships between burstiness and firing rate can also be observed in the global Pearson correlation values, which assume different values and sign in hippocampus (*r* = −0.21; *t*(18377) = −28.8, *p* < 10^−10^, Student’s *t* test), amygdala (*r* = −0.06; *t*(10148) = −6.4, *p* < 10^−10^), and entorhinal cortex (*r* = 0.11; *t*(13504) = 13.3, *p* < 10^−10^). Importantly, *LvR* displays non-trivial behaviour also locally in time, meaning that sudden changes in firing rates do not always trigger the same *LvR* variations, as displayed in Fig. 1e.

### Burstier patterns are present in all of the anatomical areas although they appear to be volatile across trials

Having established a substantial independence from firing rate values, we move now on to the specific measurements on *LvR* alone in this data set. On average, we observe a prevalence of irregular patterns (LvR = 1.19 [0.89,1.42], 1.22 [0.93,1.48], 1.19 [0.95,1.42], for hippocampus, amygdala and entorhinal cortex, respectively; mean and interquartile range (IQR) over all of the trial×unit combinations). For the sake of completeness, the maintenance and probe neurons introduced in Boran et al. (2019) were also inspected. These were identified as those with a higher spike count during the maintenance or probe period, respectively when compared to the fixation period. As shown in Fig. S1a, maintenance neurons display significantly lower *LvR* values with respect to the other units in the entorhinal cortex (*p* = 0.026, Wilcoxon rank sum test). For probe neurons, the difference depends also on the anatomical area: in the hippocampus, they are associated with lower *LvR* values (*p* = 0.040) while in the amygdala with higher ones (*p* = 0.024). Evidently, it is difficult to draw conclusions on a definite relationship between irregularities and sustained firing rates, as those shown by maintenance and probe neurons.

Next, we assessed whether the firing behaviour remains stable within a recording session by investigating the trial-to-trial variability of *LvR* of individual units, see Fig. S1b. This is examined by simply computing the mean and standard deviation of the *LvR* values separately for each unit across trials. The correlation values between standard deviation and average exhibit positive to (weakly) negative values when examining in this order hippocampus (*r*=0.573, *p* < 10^−10^), amygdala (0.225, *p* = 0.0005), and entorhinal cortex (−0.126, *p* = 0.027). Thus, in both hippocampus and amygdala, a neuron displaying burstier patterns will tend to do so inconsistently across trials whereas the opposite (to a weak extent) is observed in the entorhinal cortex.

### *LvR* values are modulated selectively by set size and trial period in the different brain regions

Next, we inspect the relationship between burstiness and the behavioural and trial variables. In preparation, we created a three-way ANOVA model between *LvR* and three factors, namely, set size, correct/wrong response, and trial period. The *LvR* was computed for each unit on 2-s windows which were centered in the respective trial periods (encoding, maintenance, retrieval; fixation was excluded due to its short duration). Since we required that each session should contain at least five trials with wrong responses (out of ∼50), we collected data from only seven sessions. The analysis delivered no significant results for the main effects, except for the trial period in the hippocampus, which is investigated later. We removed the correct/wrong response factor to enable us to perform the ANOVA test with a larger number of sessions (26, using only trials with correct responses). Trial period appeared to modulate the *LvR* response in the hippocampus (*F*_2,31502_ = 3.77, *p* = 0.023) and the entorhinal cortex (*F*_2,23340_ = 5.11, *p* = 0.006) whereas set size affected only the amygdala (*F*_2,16868_ = 9.42, *p* = 8 · 10^−5^). In Fig. 2 we show the results of the pairwise tests corresponding to those significant main effects. During maintenance periods, higher *LvR* values are observed in the entorhinal cortex and hippocampus, although the comparison was significant only with respect to the retrieval period for the latter. In the amygdala, on the other hand, we observe significantly higher *LvR* values for set size 6 when compared to both 4 and 8.

**Figure2:**
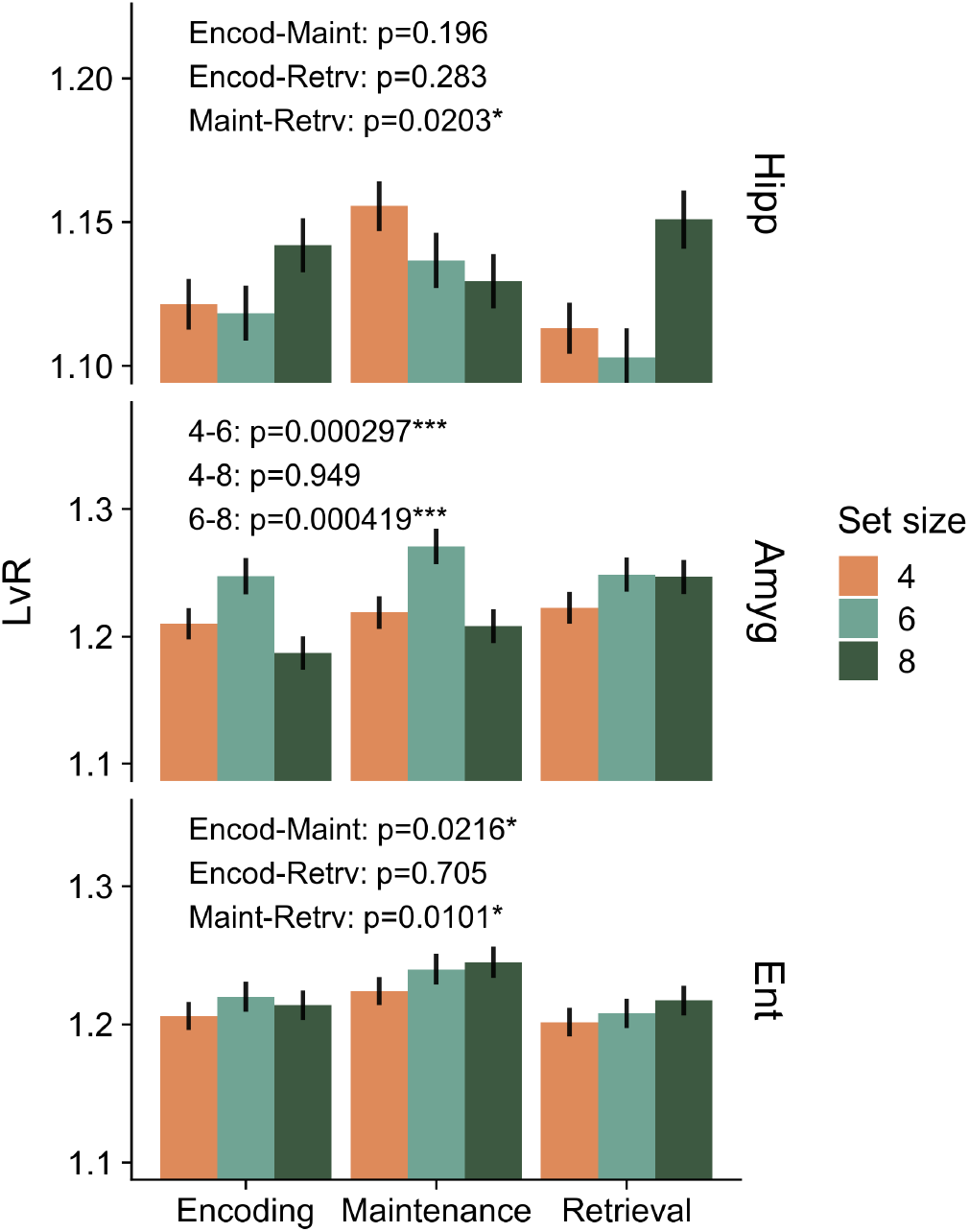
*LvR* values resolved by combination of anatomical area, set size, and trial period. *LvR* values were calculated on 2-s windows centered on the respective trial periods. A two-way ANOVA test with trial period and set size as factors was performed beforehand within each anatomical area. Only factors associated with significant main effects were examined further (interaction effects were included in the model but they did not reach significance). In particular, we show here the results (*p* values) of post-hoc pairwise *t* tests (\**p* <0.05, **<0.01,***<0.001, BH-corrected).

### 3.1 Irregularities in unit combinations and their relationship with response times

Irregularities can also occur and be monitored at the ensemble rather than the single-unit level; for this reason, we extended our analysis to include spike patterns resulting from pairs and triplets of units. In practice, for the latter, we combined the spike trains of two or three simultaneously recorded units and computed the *LvR* measure on the joint time series (see Fig. 3a). The rationale behind this procedure is that among the characterizations of a ‘persistent’ activity, the spike train of a single unit can appear sparse and irregular. However, simultaneously recorded units can fire asynchronously and together fill the activity gaps during the delay period, showing ultimately persistent and, potentially, more regular firing patterns as a whole (Lundqvist et al., 2018). On the other hand, irregular patterns can be ultimately reinforced by the combination of concomitant bursty patterns. It is thus interesting to investigate whether genuine coordination between units is present and to what extent they carry information on the trial variables.

**Figure :**
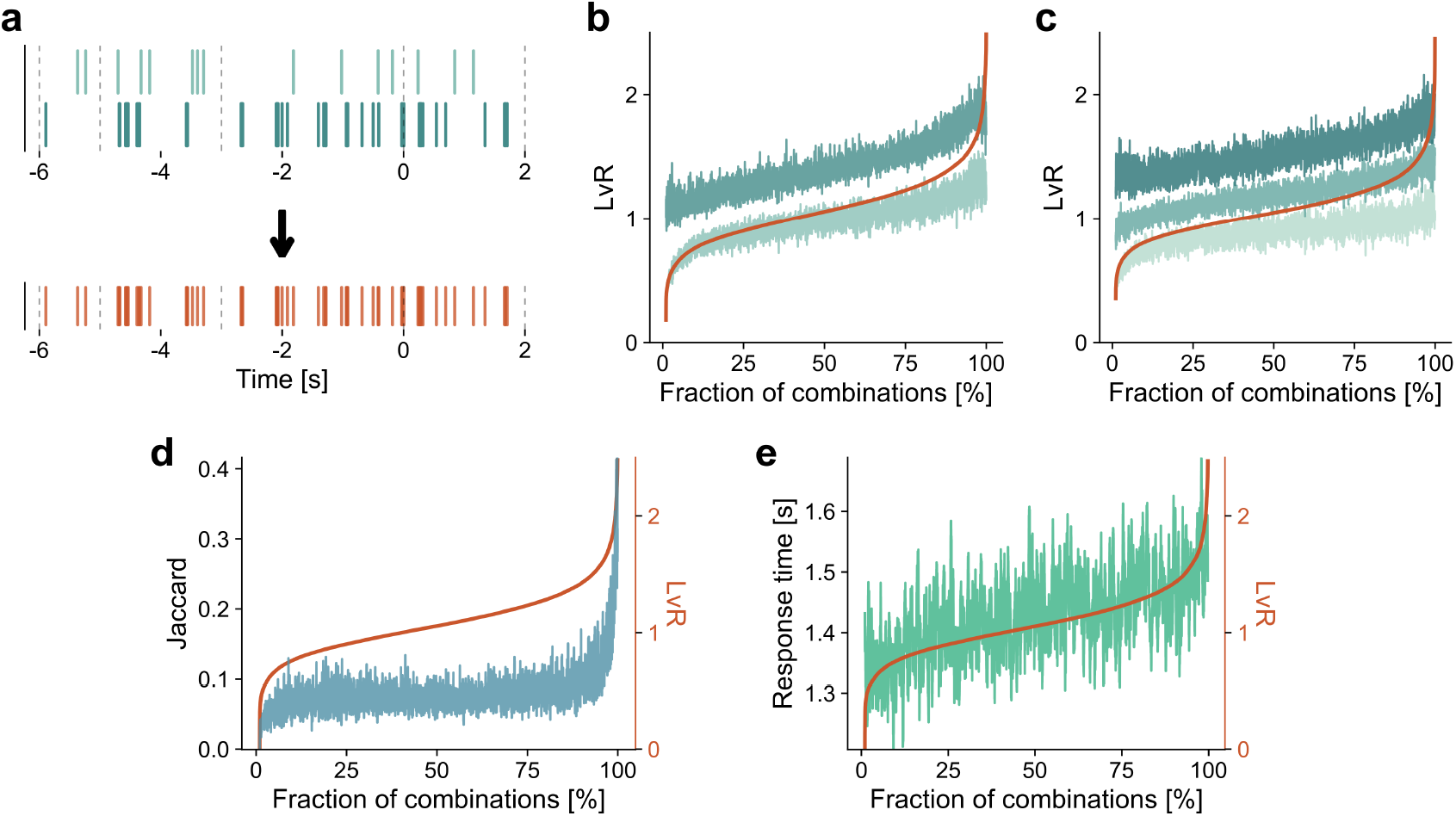
Combinations of pairs or triplets of spike trains. (a) Spike trains of two units recorded simultaneously (top) are combined into a new spike series. The dashed, vertical lines highlight the time structure of a trial (fixation, encoding, maintenance, retrieval). (b) *LvR* values resulting from the combination of two spike trains are sorted (red line) and shown along with those of their individual components (green traces). See Fig. S2 for an alternative representation of these results where the single-unit values are sorted. (c) Same as in (b) but showing *LvR* values deriving from the combinations of three spike trains. (d) Combined *LvR* values and correlations between spike trains. Jaccard values were computed between pairs of binarized spike trains (50 ms bin; blue line). These values are reordered with respect to the combined *LvR* value obtained from the corresponding unit pair (red line). (e) Combined *LvR* values and response times. Similarly to (d), the sorted sequence of paired *LvR* values (red line) is plotted but here along with the (reordered) response time of the related trials (green). In panels (b)-(e) data from all of the sessions and anatomical areas are used; not all of the combinations are plotted, but a subsample that preserve the relative contribution of each session in terms of recorded units (see below). A moving average filter with a window of 10 and 60 points is applied in panels (b)-(d) and panel (e), respectively, for visualization reasons (∼40000 points are shown)

In Figs. 3b and c we show the resultant burstiness values (red curves) combining two and three individual spike trains, respectively (green curves). The relation between the individual components’ *LvR* and the combined ones is, not surprisingly, monotonic on average (the curves shown are smoothed). The combined value remains closer to the smallest component in *LvR* (lighter hue) rather than to the highest one (darker hue) for a large fraction of the total number of combinations (around ∼80%) and *LvR* spectrum (up to ∼1.2 for the combined value). The combined *LvR* seems then to depend considerably on correlations between units in the opposite cases of strong and weak values. This is shown in Fig. 3d, where the Jaccard similarity measured derived from 50-ms bins has been adopted (not be confused with the use of the Jaccard measure described in Sec. 2.1). The lowest *LvR* values, *i*.*e*., the more regular trains, tend to be formed by uncorrelated units; on the other hand, there is a steep surge in burstiness for highly correlated neurons. This can be understood from the fact that closely coordinated spikes create smaller ISIs in the joint spike train due to overlapping bursts. This contrasts with two uncorrelated, bursty neurons, where the overlapped train is unlikely to see a strong increase in short ISIs because the burst periods do not align.

#### Burstiness shows correlations with response times independently of firing rates and with larger effect sizes for unit combinations

An interesting behavioural variable yet to be analyzed here is the response time of the patients, which provides a proxy measure for the level of attention as well as for the perceived complexity of the task. In the following, we looked for possible relationships with burstiness values. When simply ordered with respect to the *LvR* values, the response times reveal a clear increasing trend (Fig. 3e). We were curious if and how this effect varies along the time axis of the trials and if its magnitude depends on the coordination between units. To this end, we computed Pearson correlations between the response times and time-resolved *LvR* values of both single and combinations of units computed on sliding windows of 2 sec, see Fig. 4a. The significance of each value is assessed with a permutation test respecting the trial structure (transparency levels; *p* <0.05, BH-adjusted across the anatomical areas). The set of combined *LvR* values was subsampled in each session in order to match the number of units used for the single *LvR* results. In this way we guarantee a fair comparison between statistics for the different levels of combinations. More details are in Sec. 2.3.

**Figure 4:**
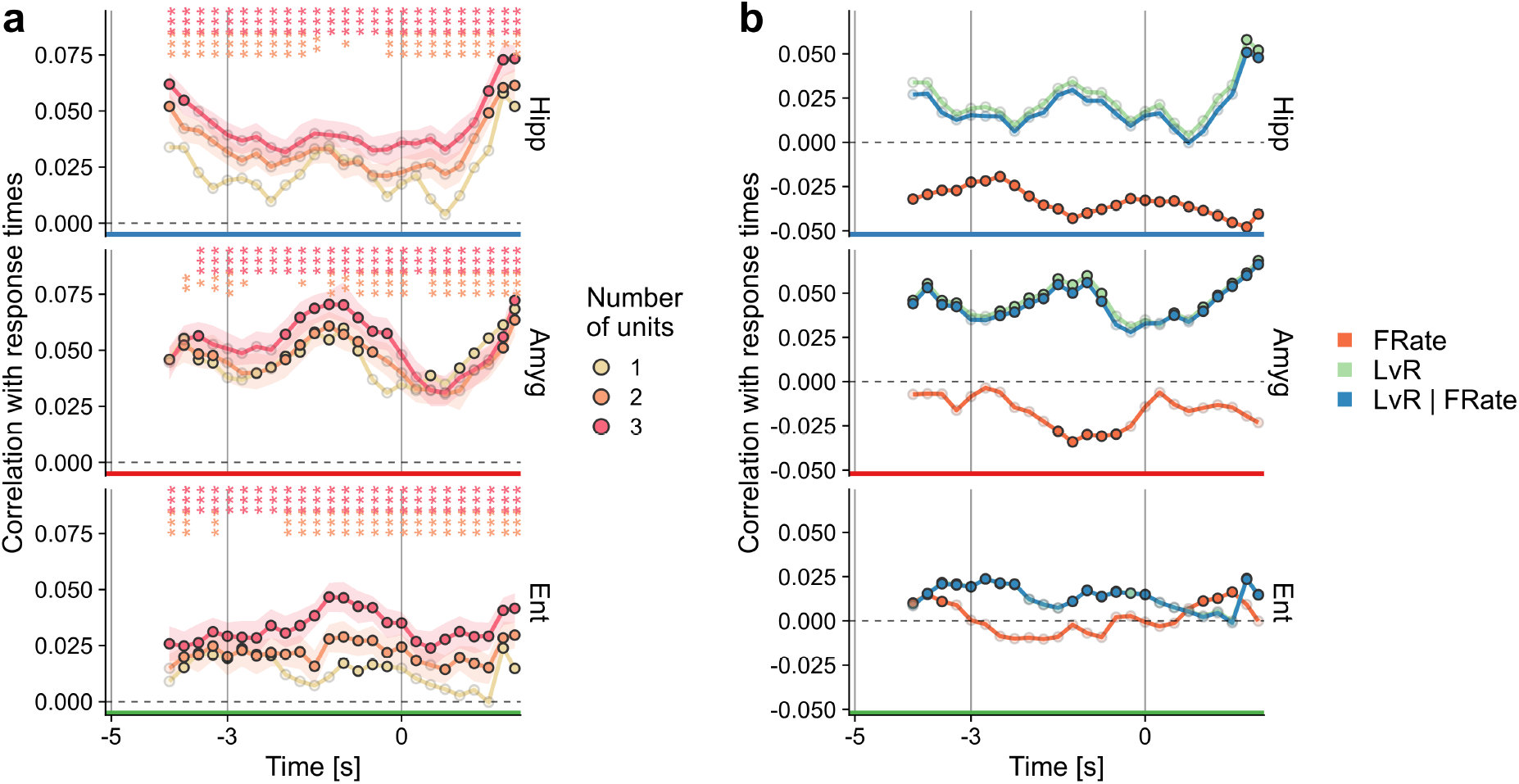
Relation between time-resolved *LvR* values and response times. (a) Pearson correlation between *LvR* and response time. *LvR* and correlation values are reported for right-aligned sliding windows of 2 s with 250 ms step (*n* = 11283, 6192, 8544 points on average for hippocampus, amygdala, and entorhinal cortex, respectively). Error bars on joint spike trains of 2 and 3 units (95% CI) derive from the repeated subsampling of unit combinations (100 times). The opacity of symbols denotes significance of the respective correlation value when compared to a null distribution (*p* value < 0.05; permutation test). Significant differences between the combined values and the single-unit ones are indicated on top (*t* test, \**p* <0.05, **<0.01,***<0.001). See Sec. 2.3 for details. The gray vertical lines delineate the encoding, maintenance, and retrieval periods. In Fig. S3 and S4, a similar approach is used for investigating the relationship between *LvR* and either set size or correct responses. (b) Contribution of firing rates. The same procedure as in panel (a) was adopted for computing the correlations between firing rates and response times. Partial correlations between *LvR* and response times, while controlling for firing rates, are shown too (‘LvR | FRate’). Only single-unit calculations are shown. See Sec. 2.3 for details.

For all of the anatomical areas, higher correlation values are observed almost everywhere for the combinations of three units (Fig. 4a). The differences between paired and single-unit *LvR* values are less pronounced in comparison. Hippocampus and amygdala show a marked modulation with respect to the specific time window. Both of the two regions reach significant Pearson correlations in encoding/fixation and in the last phase of each trial. In addition, this holds for the amygdala also for a large fraction of the maintenance period. In the entorhinal cortex, most time windows show weak but significant correlation but only if we consider combinations of two or three units. A similar time-resolved analysis was performed for the categorical variables (set size and correct response, see Fig. S3); however, we could not observe clear or significant relationships of the *LvR* values with these two variables for either single or combinations of units.

Could the correlations with response times be explained in terms of firing rates? We repeated the analysis using firing rates of single units instead of *LvR* and show the results in Fig. 4b. Generally, these latter values tend to anti-correlate with the response times with lower absolute magnitudes, except for the hippocampus where the significance threshold is met everywhere. In spite of the fact that the profiles approximately mirror those of *LvR*, the correlations between *LvR* and response times seem to be unaffected by firing rate values, as shown by a partial correlation analysis (blue line, Fig. 4b).

### 3.2 Population bursts

Next, we proceeded to examine irregular patterns at the population level. In particular, we characterized population burst events and their distributions. These events were investigated for two main reasons: first, they represent episodes which can be relevant for the underlying memory processes (Vaz et al., 2020); second, they constitute a coordinated activity between units and, thus, it might be interesting to check whether regular or irregular activities mediate this coordination. More in detail, a population burst is defined as a period where the collective activity is sustained and persists above a specific threshold computed by pooling all of the trial data, see Fig. 5a and Sec. 2.5 for details.

**Figure 5:**
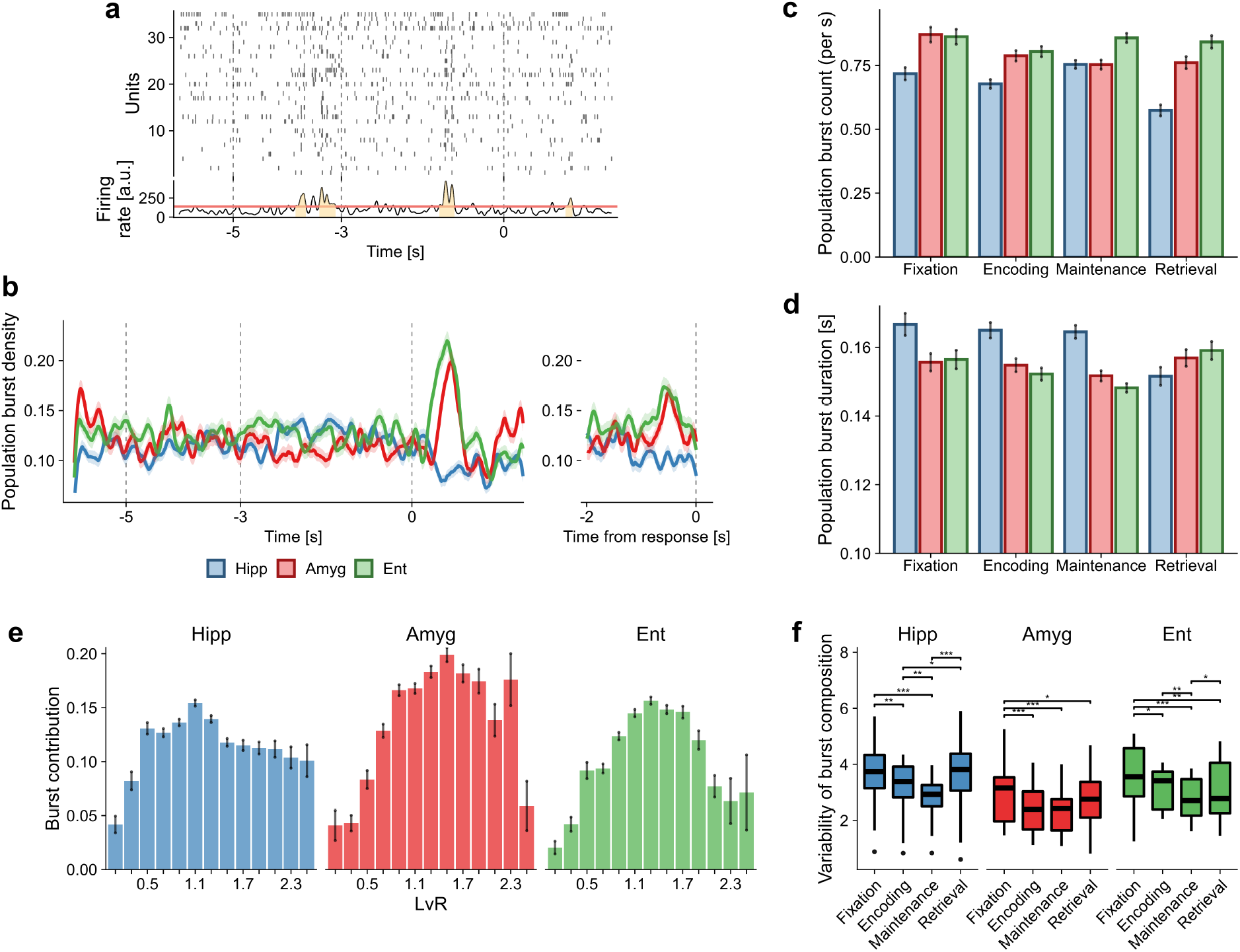
Population burst analysis. (a) Example of population burst events. Spike raster plot of an example trial (top) and the global firing rate (bottom) are shown. Burst events are defined as events longer than 100 ms where the collective firing rate exceeds a data-derived threshold. See Sec. 2.5 for details. (b) Burst density per anatomical area. The densities were calculated at each time bin (10 ms) as the fraction of trials where a burst event was ongoing. Bootstrapping at the trial level (1000 times) was used to arrive at the 95% CIs that are shown. The numbers of trials analysed were 993, 809, and 806 for hippocampus, amygdala, and entorhinal cortex, respectively. Time traces aligned with respect to the response time are shown on the right. In this case, trials with response times higher than 2 seconds were discarded (∼ 110 trials). In Fig. S5 we show the burst density profile computed excluding the probe neurons. (c) Population burst count. The number of burst events per period was counted within each trial and divided by the respective period lengths. The probe or effective retrieval period is defined only up to the response time. Mean and SEM across all of the pooled trials are shown. (d) Duration of burst events. Burst events within each trial period were pooled across all of the trials; mean and SEM of the burst lengths are shown. In (c) and (d), trials with response times higher than 2 seconds were discarded. The analysis of panels (b)-(d) where the different set sizes are distinguished is reported in Fig. S6. (e) Contribution of single units to population bursts. Each burst is characterized by specific contributions in terms of single-unit firing rates. Contributions of the single units were binned according to their underlying respective *LvR* values. For each bin, mean and SEM of the single-unit contributions to population bursts were computed. The values shown here are computed within the maintenance period; however, the distributions are qualitatively similar across the different trial periods. (f) Variability of burst composition across different trials. Distances between burst compositions across the different trials were computed. These values were then normalized with respect to those obtained from the whole trial time span (see Sec. 2.5). Results of pairwise Wilcoxon rank sum tests between the different trial periods are shown (\**p* <0.05, ** < 0.01, *** < 0.001). Further measurements regarding the bursts’ composition are provided in Figs. S7 and S8.

#### Burst activity is sharply modulated after probe presentation

Firstly, we analysed the burst density in a time-resolved manner, and this is shown in Fig. 5b, which plots the fraction of trials that display an ongoing burst event. Focusing on the hippocampus, there is a slight increase in the first two seconds of the maintenance period followed by a quick drop after the appearance of the probe letter. This behaviour is reversed for the amygdala and entorhinal cortex where an abrupt increase of burst activity within one second of probe presentation is apparent. This enhancement is likely to be triggered by the probe onset itself rather than being related to the initiation of movement or to the effective memory retrieval process. This is evident from the fact that the effective burst density is reduced if the time series are aligned to the effective response time (inset to the right), although with smaller effect for the amygdala. Can this activity be explained by the probe neurons investigated in Boran et al. (2019)? In Fig. S5, we measured the population bursts again but excluding the probe neurons from the analysis. When aligned to the probe presentation (left panels), the peak is indeed reduced in the entorhinal cortex (although still prominent) but it remains unaltered in the amygdala. On the other hand, a reduction in this area does become visible when data are aligned to the response times. Given that, we suspect that the burst activity stemming from the probe onset *per se* is likely to be evenly distributed across the population. In contrast, the signal associated with the effective memory retrieval (if any) is likely to be captured by a limited set of neurons, such as the one represented by the probe neurons.

Are these modulations related to a higher frequency of bursts or to a longer burst or both? In Fig. 5c-d we show the mean values of these quantities per trial period. For the burst frequency (panel c), hippocampal measurements tend to be lower everywhere especially with respect to the entorhinal cortex (Kruskal-Wallis test, *p* < 0.001 in all periods; post-hoc pairwise Wilcoxon rank sum tests, *p* < 0.001 everywhere except in maintenance with respect to amygdala). Only hippocampus and amygdala show differences across the periods (Kruskal-Wallis test, *p* < 0.001, = 0.045, = 0.27 for hippocampus, amygdala and entorhinal cortex respectively). The hippocampus displays the highest counts in maintenance (*p* < 0.05, Wilcoxon rank sum tests) and the lowest during retrieval (*p* < 0.001). The amygdala, on the other hand, shows a significant difference only between the fixation and the maintenance periods (*p* = 0.007). Regarding the effective average duration of the bursts, the hippocampal ones are significantly longer than those of amygdala and entorhinal cortex during the maintenance period (Kruskal-Wallis *p* < 0.001, post-hoc Wilcoxon tests all *p* < 0.01) and become shorter in the retrieval epoch when compared to the other periods (Kruskal-Wallis *p* < 0.001, post-hoc Wilcoxon tests all *p* < 0.01). Given these results, we are inclined to regard the density peaks of amygdala and entorhinal cortex seen in Fig. 5b as the result of a time-locked event confined to the first second of probe presentation, which is followed by a sustained decrease up to the response time. We will discuss this also later.

Is the occurrence of bursts or some property of bursts related to the workload? Fig. S6 shows the same analysis as Fig. 5 but additionally resolved by set size. The bursts characterizing the retrieval period are present in both of the workloads. Some interesting peaks are distinguishable for set size 4 but not for 6-8, in particular in the amygdala, but also for the other regions, a bit less than 1 s into the encoding period. No relevant and significant differences are reported for counts and duration of the bursts (Fig. S6).

#### Population bursts tend to rely more on irregular units and are unstable in composition across trial periods

Having assessed the presence of relevant population bursts during the trials, we next inspected the composition of these coordinated events. The bursts identified are characterized by the simultaneous activity of a number of units that roughly compose 50% of the population. This is shown in Fig. S7a where we plotted a sparsity measure that quantifies how much the burst firing rate is generated by a distributed activity of the units. Significant differences are observed only across anatomical areas but not across trial periods (Kruskal-Wallis test). In particular, the entorhinal cortex shows lower sparsity during maintenance than the other areas (*p* <0.001, pairwise Wilcoxon rank sum tests).

Given that not all of the units were active during the bursts, we then asked ourselves which type of neurons were contributing the most. Are both regular and bursty units active during population bursts, or is it only one type? To examine this, we binned the neurons’ activities according to their *LvR* values and computed the average burst contribution for each bin, see Fig. 5e. Across all of the anatomical areas, the most regular units (*LvR* < 0.5) do not seem to participate much in population bursts. Conversely, random or irregular units appear to contribute the most (to a similar extent), in particular for hippocampus and amygdala. Given that the distributions of Fig. 5e are approximately unimodal, it makes sense to summarize the information of Fig. 5e by extracting a single *LvR* value per burst event. Namely, we computed the mean of the single units’ *LvR* weighted by their contribution to the burst. As Fig. S7b shows, units in the hippocampus display a lower burst *LvR* than the other anatomical areas everywhere (all *p* < 0.001, Wilcoxon rank sum tests across anatomical areas), and it is lowest during the encoding epoch (all *p* < 0.05, Wilcoxon rank sum tests across trial period). Amygdala and entorhinal cortex display an increasing and a decreasing trend of burst *LvR*, respectively. However, only the results for the entorhinal cortex are significant in a Kruskal-Wallis test (*p* = 0.03) and in successive post-hoc pairwise comparisons (retrieval lower than fixation and encoding, *p* = 0.03 for both, but not than maintenance, *p* = 0.10). It is interesting also to compare the computed burst *LvR* with the single-unit values averaged within each trial (in other words, the unweighted version of the same calculation). This comparison provides further information on which type of units, on average, are more likely to participate in a burst event. Interestingly, all of the anatomical areas behave differently: hippocampal bursts tend to be composed by more regular units as evident from the fact that the weighted values are lower (Wilcoxon signed rank test, *p* values on top annotation, Fig. S7b); amygdalar ones do not systematically recruit units with specific (ir)regularities but display variability across different periods (tests are paired with significant results); finally, the population bursts in the entorhinal cortex exhibit a preference for burstier neurons (weighted values are higher).

Are always the same units constituting the bursts? Or does the composition vary substantially across the different trials? This issue is examined in Fig. 5f, where the variability of burst composition across trials has been quantified and normalized with respect to the trial-to-trial variations observed across the whole trial span (see Sec. 2.5). Positive values indicate that unit recruitment (firing rates) during bursts tends to vary more with respect to the firing rate variations across trials, and this appears to be the case for all the anatomical areas. Interestingly, there are noticeable differences between the trial periods: generally, during encoding and maintenance epochs, the composition is more stable with respect to the fixation and retrieval ones, especially for the hippocampus. Evidently, the coding of the letter set appears to induce more stable activity during windows of high firing rates (*i*.*e*., population bursts). We will investigate in the next section whether this is replicated also along the entire time span. Lastly, we check whether the unit composition of the bursts varies significantly between the different set sizes. In order to do so, we computed Mahalanobis distances between each burst and a reference set identified as those in the fixation period and then compared these distances for the different set sizes (see Sec. 2.5 for more details). Results are shown in Fig. S8. No clear differences between the workloads are visible although *p* values close to significance levels in the maintenance period of the hippocampus may suggest an increased sensibility to the neuronal coding of set size, which we discuss later.

### 3.3 Trial-to-trial variability analysis

Fano factor (FF) analysis of the variability across trials can provide further information on some of the previously presented results, for example, how does the population burst activity reflects on the FF? We start by examining the average FF values for the three anatomical areas along the trial time (Fig. 6a). The hippocampus displays generally higher variability values with respect to the other two areas (cluster-based permutation test, *p* < 0.05, Maris and Oostenveld (2007)) whereas amygdala and entorhinal cortex assume distinct values in particular during maintenance, with the former showing a more stable firing across trials. Generally, all of the areas share the same distinct patterns during the presentation of any stimulus; specifically, we recognize a drop in FF after the first second of the encoding period (∼-4 s) as well as of the retrieval period (∼1 s). For the latter, the result appears consistent with the population burst dynamics of Fig. 5b where a sustained activity of amygdala and entorhinal cortex emerged with similar timing relative to the probe presentation. On the other hand, in Fig. 6a, also the hippocampus shows a reduced variability, suggesting a consistent decrease in burst activity across trials. The inset (right panels) allows the interesting observation that, unlike hippocampus and entorhinal cortex, the FF of the amygdala is not minimal just before the time of response but rather ∼1 second earlier. Moreover, it reaches lower FF than those shown when aligned to probe presentation. Activity in the amygdala thus seems to be more locked in time to the response times (1 sec in advance); this might be indicative of the fact that the amygdala is more involved in the preparation of movement or memory retrieval than the other anatomical regions.

**Figure 6:**
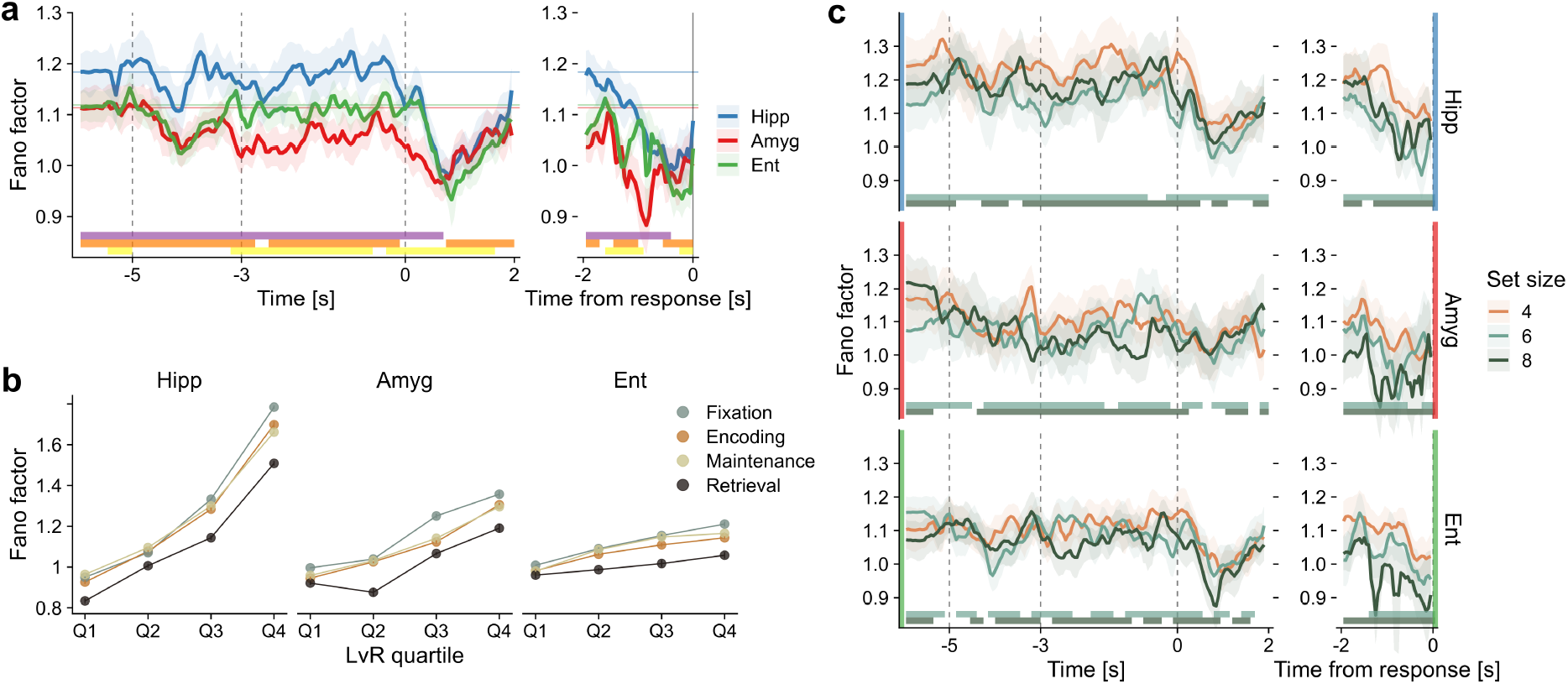
Variability of neural responses across trials. (a) The Fano factor (FF) calculation was controlled for variability in firing rates through a mean-matching procedure (see Sec. 2.4 for details). Calculations were performed on right-aligned sliding windows of 500 ms with a 50 ms time step. Mean and 95% confidence intervals are shown. Horizontal lines represent the average value of FF during the fixation period. Vertical gray lines delineate trial periods. On the right inset panel, spike trains from the different trials are instead aligned to the response time before calculating the FF. Results of a cluster-based nonparametric permutation test are shown on the bottom (*p* < 0.05; violet: ‘Hipp’ vs ‘Amyg’; orange: ‘Hipp’ vs ‘Ent’; yellow: ‘Amyg’ vs ‘Ent’). (b) The same FF calculation as in a) was repeated by splitting the neuronal population into four quartiles on the basis of their mean *LvR* values. The average FF within each trial period is reported. The results of ANOVA and pairwise tests between the different combinations of quartiles and trial periods are reported in the main text. (c) Same as (a) but distinguishing set sizes for each anatomical part. Results of a cluster-based nonparametric permutation test are shown on the bottom (*p* < 0.05). Only pairwise tests involving set size 4 are shown, with bar colors matched to the other set size involved. A moving average filter of four points (200 ms) was applied in panels (a) and (c) to improve readability.

#### Fano factor increases with burstiness values and helps in discriminating the workloads

What is the relationship of FF with the irregularities of the spike sequences? The answer to this question is not a trivial correspondence. For example, for spike trains exhibiting regular patterns, the firing rates could either vary (high FF) or remain constant (low FF) across trials. Similarly, bursty sequences could display either asynchronous patterns (high FF) or coordinated activity during specific epochs (low FF) (Constantinidis et al., 2018). If no specific time-locked activity is present, we do expect higher FF for higher *LvR* values. In Fig. 6b we split the units into four quartiles based on their mean *LvR* values and recomputed the FFs. The average values within each trial period are shown. A two-way ANOVA with trial periods and *LvR* quartiles was performed beforehand and showed significant main effects for both of the factors in all of the anatomical areas (*F*_3,481_ = [153.2, 4240.1], [110.0, 964.8], [94.48, 198.24] for hippocampus, amygdala, and entorhinal cortex, respectively; all *p* < 10^−10^. The two *F* statistics in the parentheses refer to the main effects of trial period and *LvR* quartile, respectively). The higher the burstiness (*i*.*e*., the quartile) the larger is the trial variability in virtually all of the anatomical areas and trial periods (Wilcoxon signed rank tests, all *p* < 10^−7^). Hippocampal units span a much larger spectrum of FF values when compared to units in the amygdala and the entorhinal cortex, and all of the quartiles tend to preserve roughly the same differences between the trial periods, with the retrieval epoch exhibiting the lowest FF values in all combinations of anatomical areas and quartiles (Wilcoxon rank sum tests, all *p* < 0.01).

To conclude this analysis, we investigated whether the memory workload modulates the trial-to-trial variability. When discriminating set sizes (Fig. 6c), we observe significant differences in FF values in all of the trial periods, including fixation. Whether this is due to differences in the number of trials per set size (there are generally more trials with set size 4) is arguable, as the relative order of the FF in the different workloads varies across the anatomical areas, *e*.*g*., during fixation. Set size 4 appears to be significantly higher than the others for the first two seconds of the maintenance period in hippocampus and amygdala and also during retrieval when aligned to response time (right panels). This suggests that higher workloads require more stable activity for both memory phases although this seems to loose stringency at the end of maintenance, possibly due to difficulties in retaining the stimulus content. We observe also that the anticipation of minimal FF with respect to the response time of the amygdala is only the result of higher workloads (6 and 8). Given that, we conjecture that this signal involves processes related to mnemonic functions rather than movement ones. During encoding, however, it is not possible to associate the (weaker) FF decrease on the first stimuli presentation with a specific workload (Fig. 6c, main panels).

### 3.4 Decoding analysis

In the final part of our analysis, we asked ourselves whether the decoding capabilities of the population activity observed in Boran et al. (2019) depend on their firing irregularities. In particular, is the decoding of the workload dependent on the *LvR* values of the underlying populations? In order to assess the importance of the firing behaviour, we split again the neuronal pseudo-population, obtained by pooling all of the sessions, into four quartiles defined by the average *LvR* value of the units. For each of these sub-populations, a linear decoder was trained and tested following a cross-validation procedure. In order to avoid confusion with the analysis of Fig. S3a, we remark that the decoding is performed in an N-dimensional space, where N represents the number of neurons, using the (normalized) firing rates. Overall, we follow the analysis performed in Boran et al. (2019) but with two main differences: we use here a decoding scheme based on bootstrap for the alignment of trials belonging to different sessions, and, as noticed we distinguish the different units according to their burstiness typology (see Sec. 2.6 for more details).

#### Neural populations showing non-Poisson firing predict the memory workload more accurately

We show the results of the analysis for the maintenance period in Fig. 7a. We observe that encoding of set size generally benefits from all of the anatomical areas and from the presence of all of the neurons irrespective of their burstiness type. However, when grouping the neuronal populations by their *LvR* values, some differences arise between the anatomical areas. Units in the hippocampus show significant decoding accuracies for all of the quartiles except for the second one (corresponding to *LvR* values between 1.03 and 1.16), with the most irregular neurons performing best. A decreasing trend in performance is observed for the amygdala whereas units in the entorhinal cortex approach the significance threshold only on the last quartile (*LvR* > 1.30, *p* = 0.065) encompassing the most irregular neurons.

**Figure 7:**
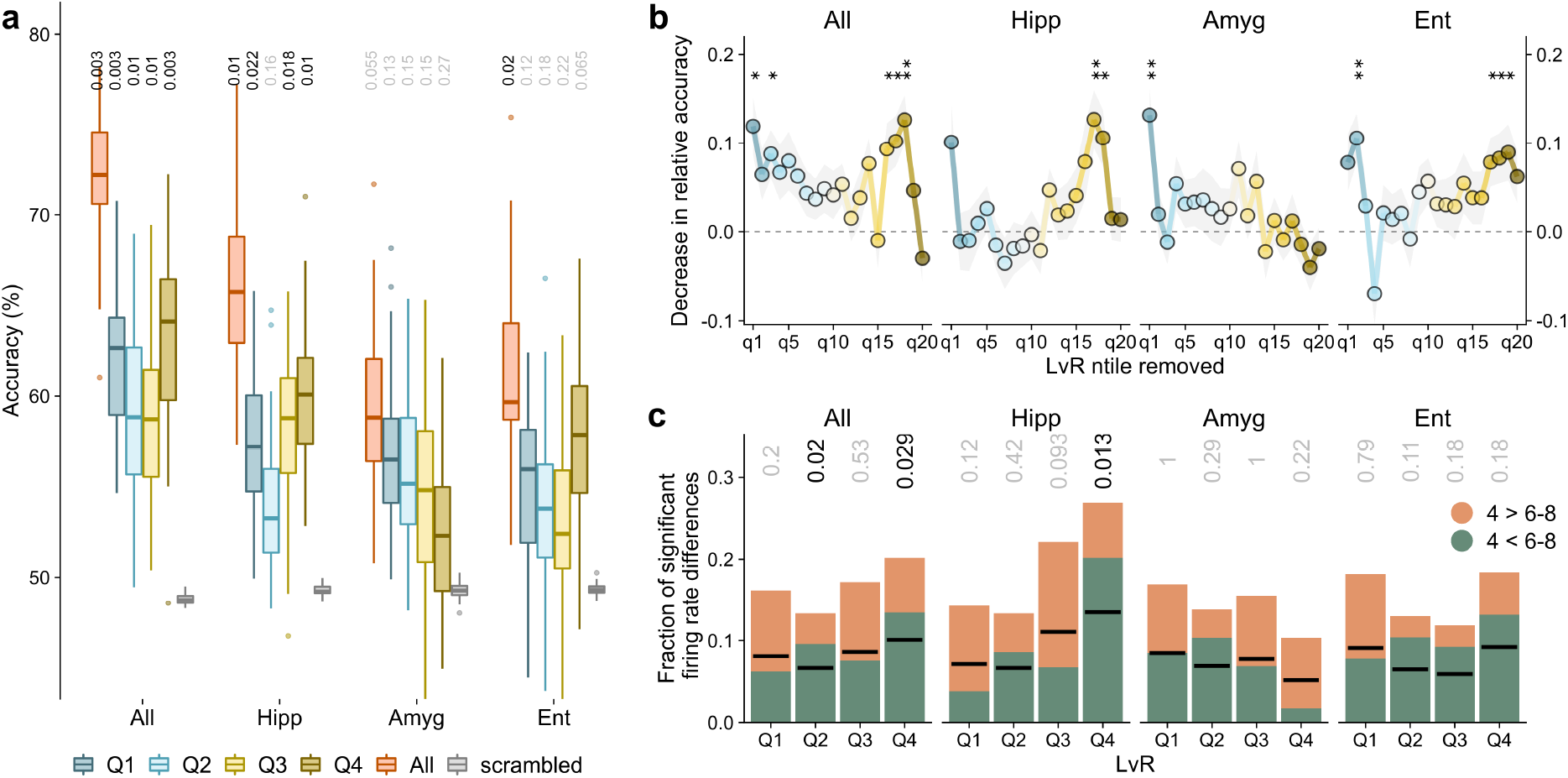
Decoding of trial variables during the maintenance period for different firing patterns. (a) Decoding performances of set size (left) during the maintenance period are evaluated on neuronal subpopulations split into four quartiles according to their *LvR* measure (blue to yellow boxplots). Decoding accuracies for the whole set of units are shown in red along with chance values obtained by shuffling the trial variable labels (grey). The boxplots represent the distribution of 50 accuracy scores obtained by a bootstrap process aimed at sampling different alignment configurations between trials of different sessions. Within each bootstrap cycle, a statistical test is performed using the chance level values obtained but scrambling the labels (500 times). The 50 resultant *p* values are summarized in the median value shown on top of each boxplot (BH-corrected within each anatomical area). We employed a balanced linear SVM model trained and tested with a 10-fold cross validation procedure. See Sec. 2.6 for more details. (b) Contributions of *LvR* sub-populations to the decoding performances. The higher the loss in accuracy (*y* axis) the larger is the contribution of the specific *LvR* n-tile to the decoding (*x* axis). Similar to a), neuronal populations are split into 20 n-tiles according to the their *LvR* values, and, in turn, each n-tile is removed and the decoding procedure repeated as above. Here we capture the normalized decrease in accuracy performances with respect to the full population scores (see Sec. 2.5). Mean and SEM of the indexes computed across the bootstrap runs are shown. Results of a *t* test with zero as expected sample mean are reported on top (*df* =49, \**p* <0.05, **<0.01, ***<0.001; BH-corrected across the different n-tiles). In Fig. S9 the same analysis of panels (a)-(b) is performed for the decoding of correct/wrong responses. (c) Differences of single-unit firing rates between different set sizes. For each unit, a *t* test between the firing rates of trials with set sizes 4 and 6-8 is performed (data points are the same as those utilized for panels (a)-(b)). The fraction of units yielding a significant test is reported while distinguishing also the sign of the *t* statistic. Black segments are positioned at half height of the respective columns to guide the eye. Binomial tests on the number of occurrences of significant positive (or negative) *t* statistics were performed (*p* values shown on top).

We switched then to a complementary approach to investigate at finer resolution how firing characteristics affect the decoding capabilities. Rather than splitting the whole population into non-overlapping sub-populations, we remove in turn a small set of neurons equal to 1/20 of the whole number characterized. This small set, in the same spirit as before, corresponds to a specific *LvR* n-tile. The resultant decoding accuracies are compared to the full population ones, which allows a fairer comparison between the two measures in terms of dimensionality of the decoding space (differing by only 1/20 and not by 3/4 as before). In more detail, we compute sensibility indexes between the accuracy scores of full and depleted populations while accounting for the variability between the cross-validation folds (see Sec. 2.6). The results are shown in Fig. 7b. Qualitatively, the same patterns reported in panel (a) concerning the set size decoding emerge also at finer resolution. However, wider variations between contiguous n-tiles are resolved, especially close to boundary values. For example, in the hippocampus, the very last n-tiles do not contribute significantly, and generally, across all of the anatomical areas, neurons firing with regular patterns contribute only with the lowest *LvR* values (first one or two n-tiles). Importantly, and this holds for all of the anatomical areas, the random (Poisson) patterns seem not to carry any relevant information about the workload. We applied the same procedure of Fig. 7 for decoding correct and wrong responses from the population activity during maintenance (Fig. S9). Somewhat surprisingly, not a single set of units reported a significant accuracy.

#### Single-unit activity offers insights into mechanisms of workload decoding

In the final part of our analysis, we tried to shed light on the previous decoding results by investigating how the single-unit firing rates coded for workloads in the different *LvR* quartiles. In Fig. 7c we plot the fraction of single units which displayed significant differences in firing rates between the set sizes (4 vs 6-8 as for panel (a)). Qualitatively, the results across the quartiles follow the same trends as observed in Fig. 7a, hinting at the fact that much of the workload information is stored in single-unit activity. Given this, it is interesting to observe that for the most irregular and significant quartiles of Fig. 7a (that is, for ‘All’ and ‘Hipp’), the number of units responding for large set sizes (6-8) is significantly higher than that favouring lower workloads (binomial test, Fig. 7c, top). We therefore conjecture that burstier units tend to possess a simpler code for the workloads, which is based on a coordinated increase in firing rates rather than on an asynchronous mixed activity across units.

## 4 Discussion

The discharge patterns that are observed in the MTL during different memory phases are hypothesized to contain information about the memory items as well as shed light on the underlying neuronal dynamics that govern the memory processes. Differently from the approaches adopted in many prior works (Kamiński et al., 2017; Boran et al., 2019; Kornblith et al., 2017; Kamiński et al., 2020; Derner et al., 2020; Bausch et al., 2021) (see also Rutishauser et al. (2021)), we did not, except for controls following Boran et al. (2019), distinguish neurons based on their firing rate levels associated with external variables and observables, such as trial periods, task conditions, or memory items. Such an approach contains the implicit assumption that only a small fraction of neurons carry information about behaviour and stimuli. Instead, we kept our focus on the whole set of recorded neurons and analysed how the (ir)regularity levels at different resolution stages, *i*.*e*., single-unit, unit combinations, and population level, appear to carry information about the different trial variables. Furthermore, we moved away from a spike rate-centric approach and from an analysis driven by the external variables and instead focused on the internal activity (Buzsá ki, 2020). The latter analysis has higher exploratory power as it is able to assess also WM models that differ from the ones built on the hypothesis of persistent neuronal activity (Kamiński and Rutishauser, 2020).

### Burstiness levels are indicative of response times and attention levels

Our analysis is based in large part on the quantification of the irregularities of spike trains through the *LvR* metric, which has a non-trivial albeit weak interdependence with net firing rates (Shinomoto et al., 2009) (Fig. 1d-f). We found that higher *LvR* values and, thus, burstier patterns tend to be associated with higher response times, with particular emphasis for the entorhinal cortex and the amygdala during the delay (maintenance) period. Irregularity patterns resulting from the combinations of multiple units (two and three) were generally more correlated with observed response times, suggesting that ensemble-level coordination between units can better encode behavioural responses. Moreover, since burstier patterns, which are associated with slower responses, are related to higher correlations between units (Fig. 3d), it would be interesting to explore whether they play a role in describing/controlling attention levels (as in Cohen and Maunsell (2009)). The fact that significant correlations are observed in the beginning of the encoding period, comprising also fixation, suggests that the attention level should be considered when interpreting the results obtained. The relation between attention and WM is not yet clear, both on the behavioural level but especially in its realization in the neural substrates (Shevlin, 2020; Oberauer, 2019). Our results hint at the discharge patterns as a variable of interest to disentangle the aforementioned relationship. For example, specific memory activity might be mediated by spike counts of the maintenance neurons of Boran et al. (2019) or more generally through memory-selective cells (Rutishauser et al., 2021) while the nature of the discharge patterns might relate only to attention-related variables. Indeed, we did not find any relevant workload dependence of the *LvR* values in the MTL regions, except for a puzzling preference in the amygdala of high burstiness values for the intermediate set size 6 (Fig. 2).

### Persistent vs. transient activity: potential integration of the two models

Considering the discussions around the competing theories underlying WM dynamics (mostly) in the prefrontal cortex (Constantinidis et al., 2018; Lundqvist et al., 2018; Masse et al., 2020), comparatively less research on this topic has been performed relying on data from the MTL areas analyzed here (Kornblith et al., 2017; Kamiński et al., 2017; Boran et al., 2019). The main discussion between the prevalent models postulating asynchronous, persistent activity against those which embrace a more transient (bursty) but coordinated (or also silent) activities is hampered by the definition of ‘persistent’ activity. This is a term that is adopted widely in the literature but, as observed in Kamiński and Rutishauser (2020), it frequently remains ambiguous and lacks a sound quantitative definition. For example, it is not always clear which neural substrates and at which resolution this activity should be observed and defined as such, *i*.*e*., single neuron (more often), population level and local networks, or brain oscillation (Leavitt et al., 2017). In addition to that, it must be remarked that each brain area can show intrinsically diverse activity characteristics during the delay period; this activity, in turn, can depend also on the stimulus type and on the structure of the memory tasks (Leavitt et al., 2017; Christophel et al., 2017; Sreenivasan and D’Esposito, 2019). Here, we choose to equate persistence with the regularity (or randomness) of the spike trains and the competing transient activity with the burstier patterns (high *LvR*). However, for the aforementioned reasons, it is important to always interpret this terminology with caution.

Our analysis shows that the successful retrieval of information in memory (correctness of the response) and the workload (set size) cannot be decoded with clarity from the spectrum of burstiness values of single units or combinations thereof (two or three) during the maintenance period (Figs. 2 and S3). Similarly, the frequency and properties of population bursts were not particularly helpful toward this goal either (Fig. S6). On the other hand, when decoding the workload from the (pseudo-)population activity patterns, our results suggest that in the hippocampus and, to a lesser extent, in the entorhinal cortex, both neurons with regular and with bursty patterns concomitantly provide a significant contribution toward decoding performance with the latter showing slightly larger accuracies (Fig. 7a-b) when their respective contributions are isolated. Possibly, the burstier the neurons the more linear is the relationship between firing rates and set size, favouring thus a better discrimination of workloads already at the single-unit level. This would be consistent with the data for the hippocampus in Fig. 7c where the quartile of highest burstiness is the only single-region one returning a significant result on the binomial test measuring preference for which set size corresponds to higher firing rates. In contrast to the other two regions, neurons in the amygdala tend to exhibit better performances the more regular their spiking activity is. Our results thus do not exclude the possibility of an integration of the two aforementioned competing models (transient and persistent) within the same brain system. However, as mentioned above, we cannot and should not rule out the possibility that one signal corresponds to the actual memory content while the other encodes the level of general attention that can certainly be modulated by the memory load.

In a recent work (Li et al., 2021), it was shown that computational models of WM implementing either burst-coding or elevated persistent activity could be distinguished by measurements of trial-to-trial variability. In particular, for the first class of models (‘burst-coding’), the authors predicted an increased FF during the delay period while the second class (‘elevated persistent’) displayed more stability across trials (lower FF). Our own results do not favor either model clearly. We did not in fact observe variations in FF between the fixation and the maintenance periods (Fig. 6a), except for a decrease in the amygdala that actually stretches into the encoding period. The same observation emerged when distinguishing neuronal sub-populations by their burstiness as characterized by *LvR* quartiles (Fig. 6b).

We did observe an important modulation of the population burst activity subsequent to the probe presentation (∼0.5 s after; Fig. 5). The enhanced activity in the entorhinal cortex and amygdala at population level is consistent with the identification of ‘probe’ neurons at single-unit level in Kamiński et al. (2017); Boran et al. (2019). However, this specific neuronal set is not entirely accountable for these peaks of activity, as these persist, at least partially, even in the absence of probe neurons (Fig. S5). As suggested in the cited works, these activity signals likely indicate a switch between a memory maintenance phase and a retrieval process (Kamiński et al., 2017). This applies especially to the entorhinal cortex. For the amygdala, we observe a minimum in FF that is time-locked to the response time (1 sec before) and modulated by higher workloads, which suggests a possible involvement in memory retrieval functions (Fig. 6a,c). The net activity decrease observed in the hippocampal population is however less clear.

### Final remarks

Summarizing, three main results emerge from the analysis of burstiness, which is a measure of the irregularity of the spike trains, during a WM task in the MTL. First, the promptness of the response, but not the workload, is inversely correlated with burstiness levels (Fig. 4a). The occurrence of these correlations already in the early phases of the task indicates a possible dependence on the global level of attention in the test subjects. Second, probe presentation is characterized by different degrees of burst activity at the population level: strongly enhanced in the entorhinal cortex and amygdala but suppressed in the hippocampus (Fig. 5). These signals are distributed across the whole population of neurons and may signify a retrieval of information from memory for the amygdala. Third, firing rates associated with non-Poisson (non-random) firing, either regular or bursty, can predict better the memory workload than those related to random patterns (Fig. 7). Our results suggest that WM might be maintained through an interplay of heterogeneous spiking dynamics, without clearly favouring either of the proposed models (persistent vs transient).

The activities of the regions analysed here (hippocampus, amygdala, and entorhinal cortex) show distinct features in the spiking patterns and in the association to behavioral and trial data. Overall, they indicate that a broader perspective of the timing and structure of spiking patterns, not constrained to the concept of persistent (enhanced) activity, is needed. Future studies could examine if interactions between these brain areas also carry information about WM performance. In addition, they could unveil whether and how the different classes of firing patterns (regular, random, bursty) are generated internally or whether they are induced from a neighboring area (Sreenivasan and D’Esposito, 2019). It is likely that heterogeneous references will be more adequate than a simple, standard WM experiment for investigating the discharge patterns. An increased complexity of the tasks, with, *e*.*g*., variable length delays, distracting stimuli or different probing schemes (Bausch et al., 2021) but also extended fixation periods could offer more insights into the stability of the spiking sequences.

## Acknowledgements

The authors thank Davide Garolini for interesting discussions and Johannes Sarnthein for having publicly shared the data set and for valuable suggestions and clarifications. This work was supported financially by an excellence grant of the Swiss National Science Foundation (310030B 189363) to A.C.

