## Supplementary figures (S1-S9) for "Spiking burstiness and working memory in the human medial temporal lobe"

### Supplementary data

Francesco Cocina[1], Andreas Vitalis[1*], and Amedeo Caflisch[1*]

[1] *Biochemistry Department, University of Zurich, Zurich, Switzerland*

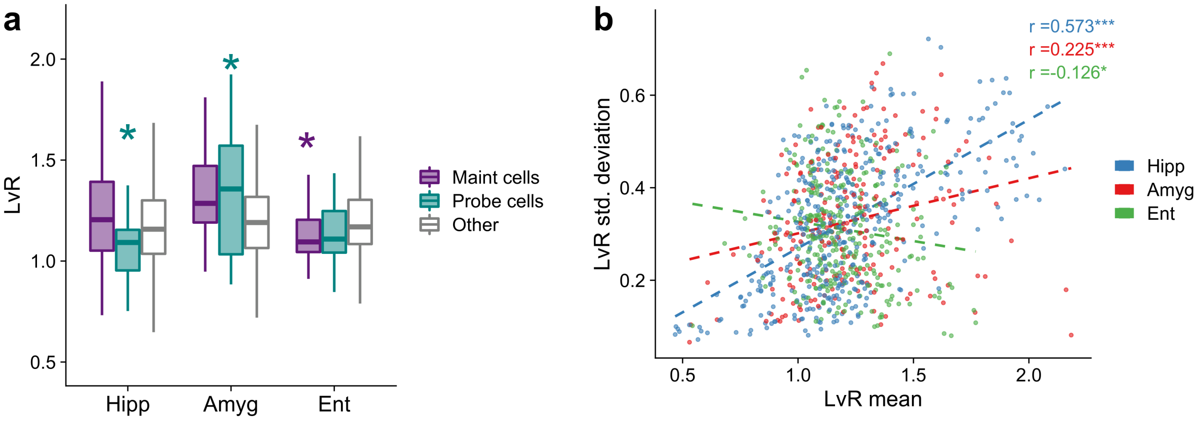


Figure S1: (a) $LvR$ values for maintenance and probe neurons. These were identified following the same procedure as Boran et al. (2019). They are characterized by a significant increase of spike counts, relative to the fixation period, in the maintenance and probe periods, respectively. A Wilcoxon rank sum test is performed for both sets of neurons against the remaining units that were not specifically characterized ('Others'). (b) Trial variability of $LvR$ values. Scatter plot of mean and standard deviation computed across trials for each unit. A linear fit is plotted to aid visualization. Pearson correlation values are shown on top with significance levels (Student's $t$ test, $t(416; 231; 304) = (14.2; 3.51; -2.22)$). $*p < 0.05$, $** < 0.01$, $*** < 0.001$.

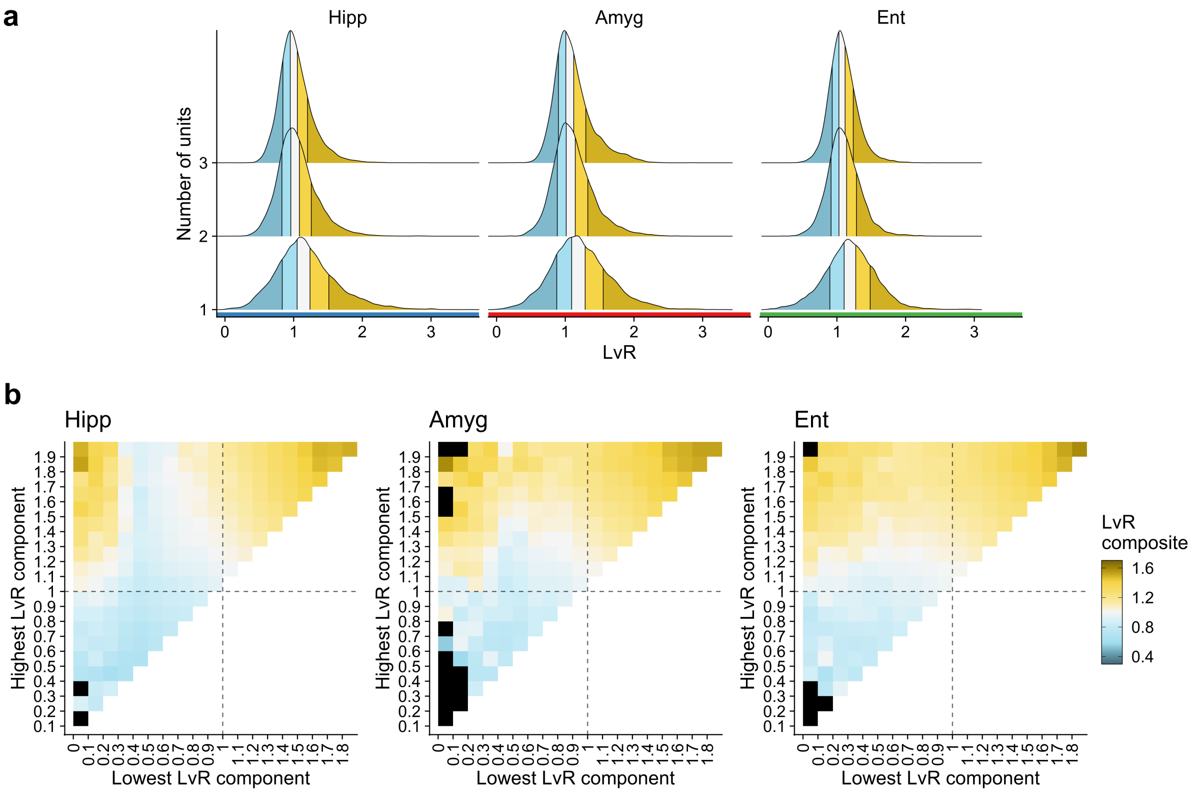


Figure S2: (a) Density distribution of $LvR$ for single units ("1" on $y$ axis) and combinations of these ("2" and "3"). For the latter, similarly to the results of Fig. 3, we randomly sampled in each session a number of points equal to the number of recorded units. This was done in order to avoid that sessions with the largest number of neurons dominated the overall sample (as the number of combinations increases with the binomial coefficient). Colors indicate the five n-tiles. (b) $LvR$ values resulting from the combination of two spike trains. The $x$ and $y$ axes indicate the values of the constituent single-unit $LvR$ values (threshold at 2); each composite $LvR$ shown is taken as the mean of the values appearing in that particular bin. If the samples are fewer than 5, the average is not computed (black bins). Analogously to the considerations done in panel (a), the averages are performed by weighting such that each session contributes with the effective number of recorded units to the final result rather than with the total number of combinations.

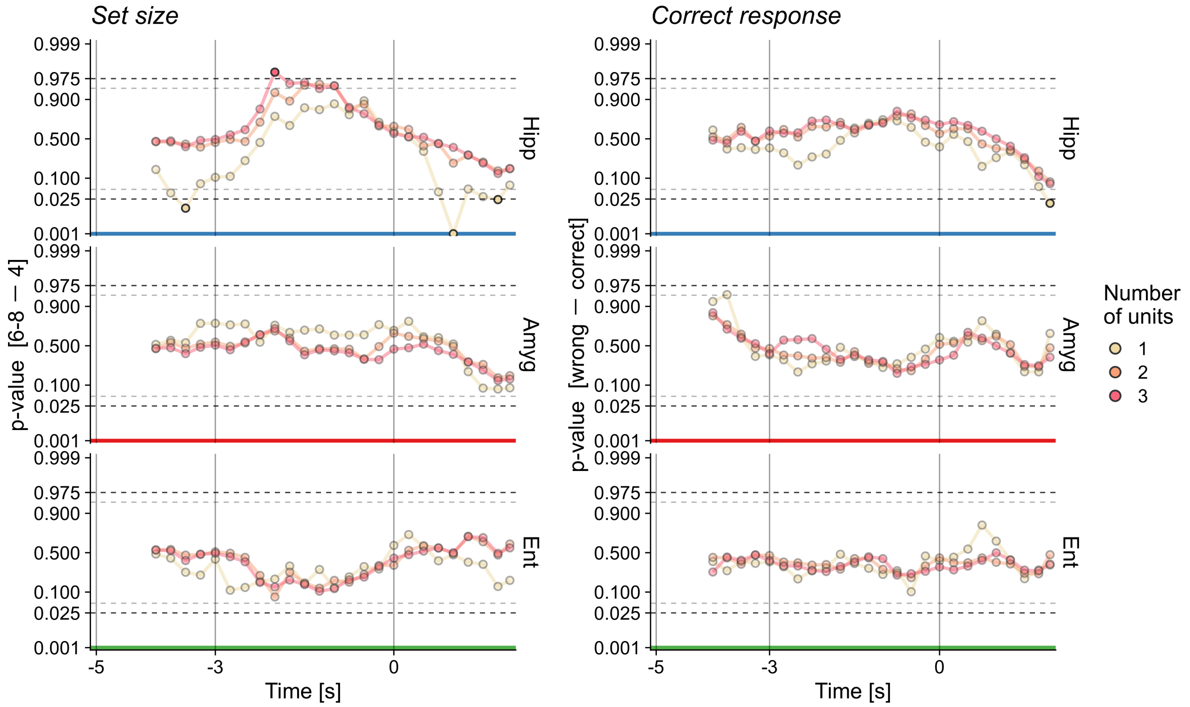


Figure S3: (a) Dependency between set size and burstiness spectrum. To quantify differences between the $LvR$ distributions of set size 6-8 *vs.* 4, we computed a test statistic that measures the signed area between the two cumulative distribution functions, similar to the Wasserstein distance. The calculation follows the same sliding window approach as in Fig. 4. The $p$ values derive from a null distribution generated by shuffling the set size labels but preserving the trial structure; the opacity of symbols denotes significance (dotted lines as two-tailed thresholds corresponding to $p$ value $< 0.05$). See Fig. S4 for the underlying test statistic values. A probit transformation is applied to the $y$ axis to widen the intervals of significance. (b) Same procedure as in panel (a), but distinguishing correct and wrong responses rather than set sizes.

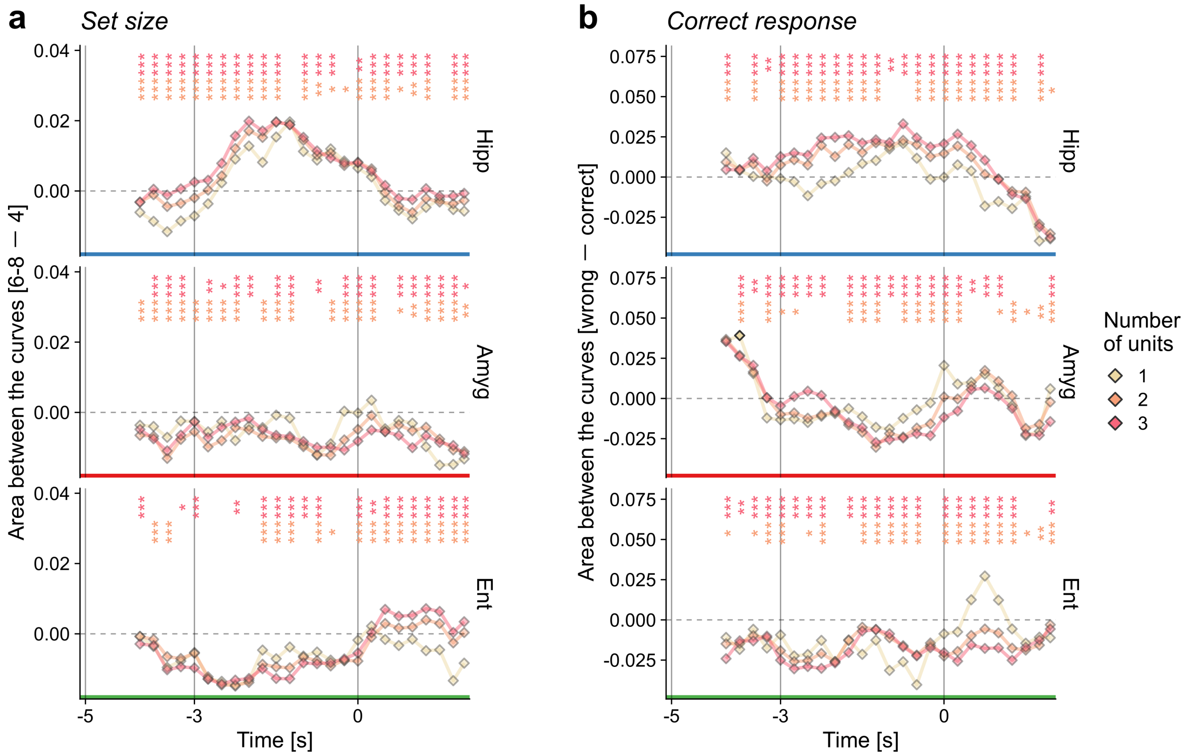


Figure S4: In panels (a) and (b) we show the test statistic values associated with the $p$ values displayed in Figs. S3a-b, respectively. The opacity of the points indicates the significance of single measurements (equivalent to Fig. 4). For each time point, the annotation on top denotes the significance level reached by a $t$ test between the combined measures ("2" or "3", 100 samples each generated by bootstrap, see Sec. 2.3) and the single-unit value, taken as the reference mean of the null hypothesis (*$p$ <0.05, **<0.01, ***<0.001, BH-corrected).

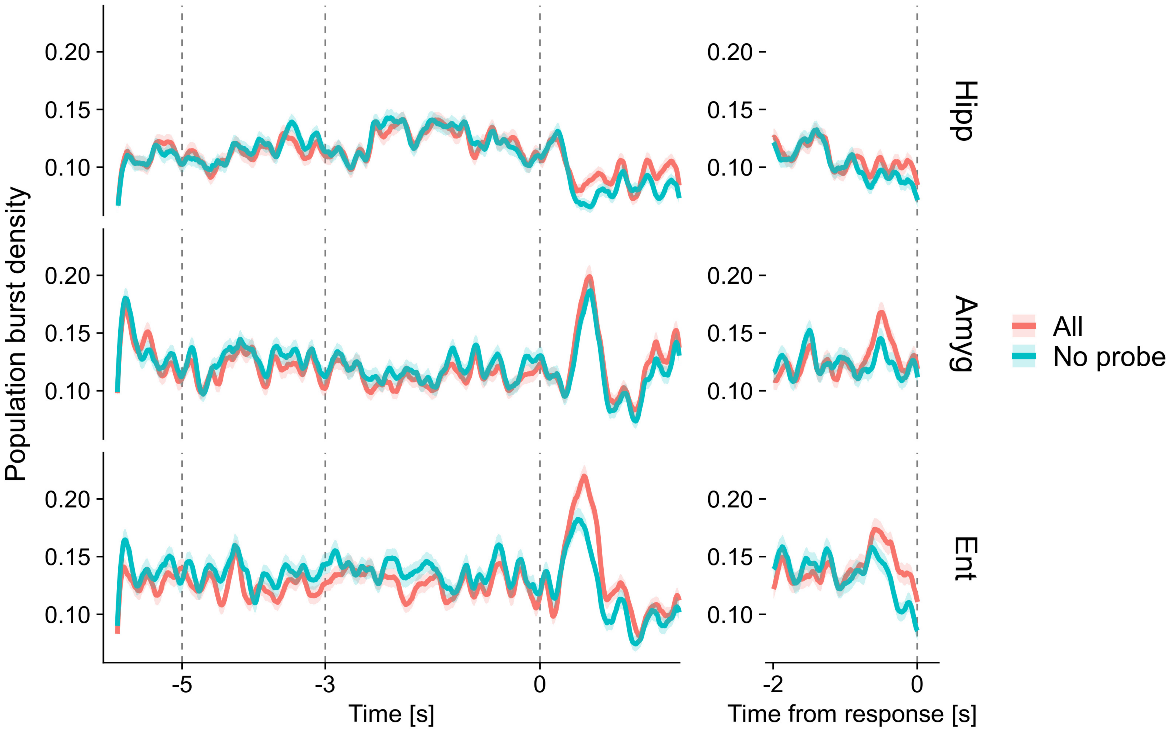


Figure S5: Density of population bursts without probe neurons. Bursts were identified excluding the probe neurons from the neuronal population ('No probe'). The profiles labelled as 'All' are the same as in Fig. 5b.

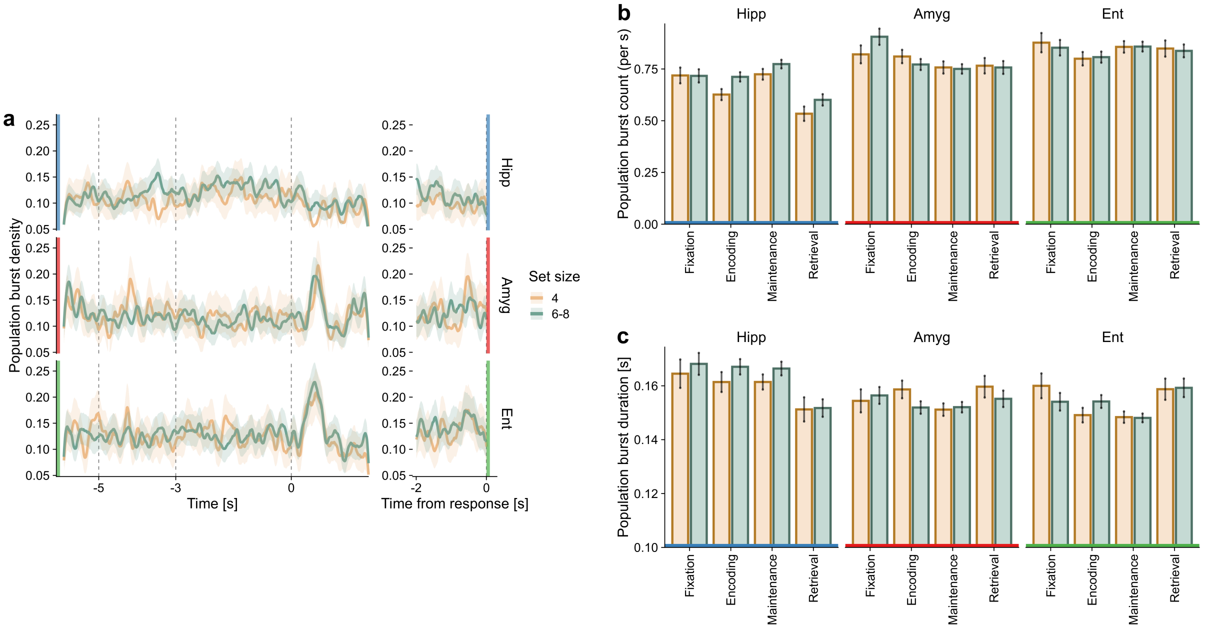


Figure S6: Population burst results for different set sizes. The analyses shown in panels (a)-(c) correspond to those presented in Figs. 5b-d, respectively.

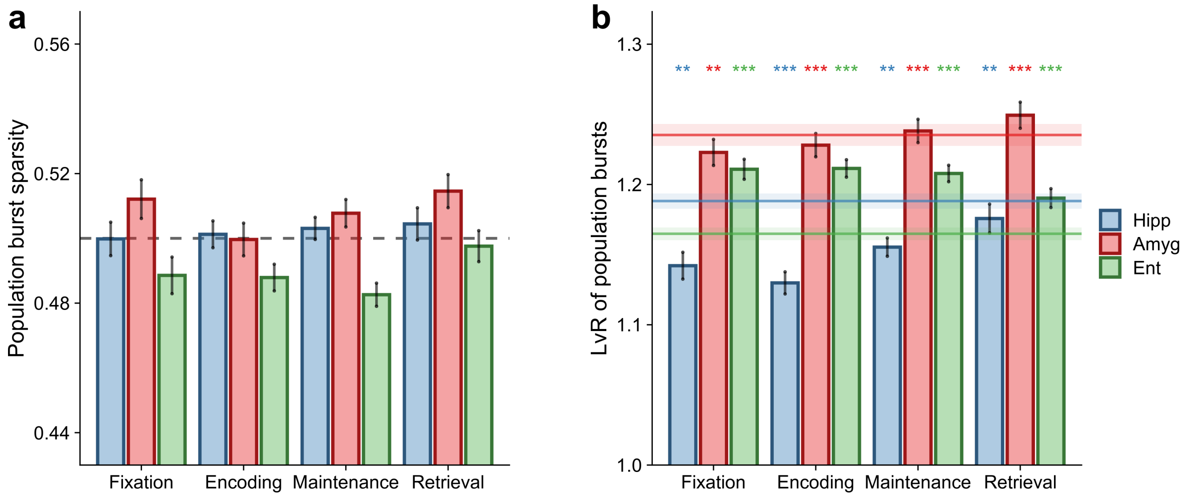


Figure S7: Properties of population bursts. (a) Sparsity of the burst unit components. Sparsity values quantify the degree of contribution of unit activities to the overall firing rate during bursts. Low values indicate that only few units are active during the burst whereas higher values point to a balanced contribution of a large fraction of the units (see details in Sec. 2.5). We computed averages within each trial and report here the mean and SEM across the whole pool of trials. (b) Burstiness of the contributing units. We computed for each burst a $LvR$ value by taking the weighted mean of single-unit $LvR$ values. The weights were proportional to the firing rates during the burst event window (unit contributions, see details in Sec. 2.5). Horizontal lines indicate the average $LvR$ values of single units. The plotted values are the mean and SEM of trial-averaged $LvR$ values of bursts computed. Wilcoxon signed rank tests were performed between burst and single-unit results for each combination of anatomical area and trial period ($n$=680, 607, 632 average number of trials for hippocampus, amygdala, and entorhinal cortex, respectively; *$p$ <0.05, **<0.01, ***<0.001, BH-corrected).

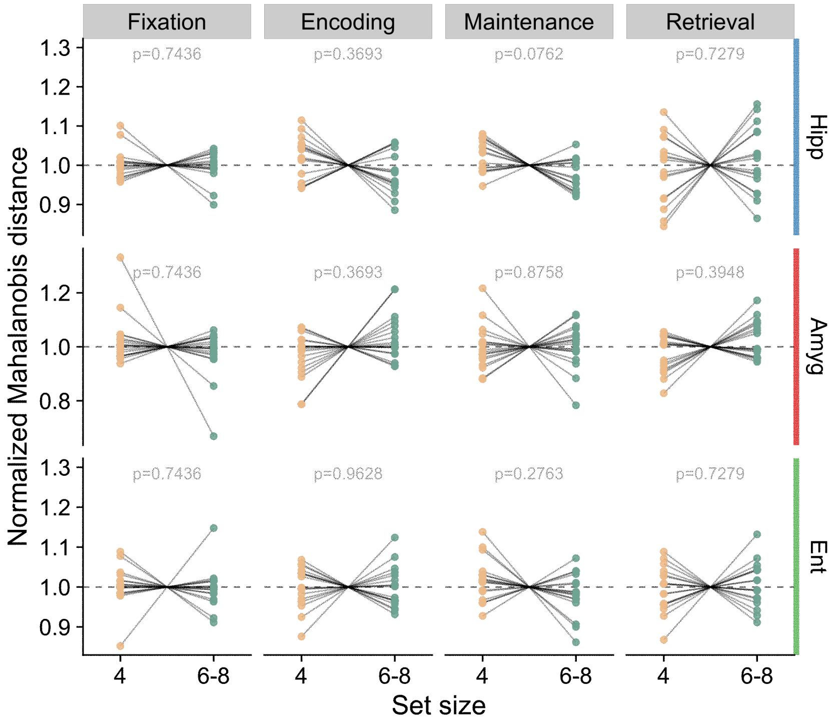


Figure S8: Distance between burst compositions of different set sizes. Within each session, Mahalanobis distances were computed between single bursts and those in the fixation period (reference group). The chosen metric space are the mean unit activities during the bursts (see details in Sec. 2.5). Average values of Mahalanobis distances were obtained for both of the set size classes and, afterwards, normalized by the sum of the two. $p$ values are shown on top of each sub-panel (Wilcoxon signed rank or $t$ tests, $n = 16$, 18 and 17 for hippocampus, amygdala, and entorhinal cortex, respectively; BH correction across anatomical areas).

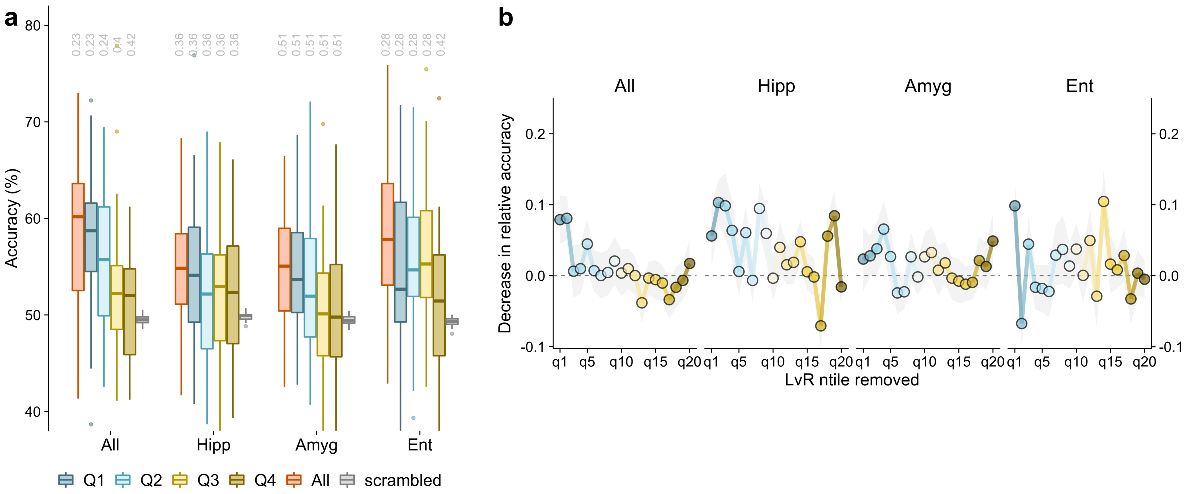


Figure S9: Decoding of correct response from population activity during maintenance. The analyses shown in panels (a)-(b) correspond to those presented in Figs. 7a-b, respectively.
